# The cytokine receptor Fn14 is a molecular brake on neuronal activity that mediates circadian function *in vivo*

**DOI:** 10.1101/2024.04.02.587786

**Authors:** Austin Ferro, Anosha Arshad, Leah Boyd, Tess Stanley, Adrian Berisha, Uma Vrudhula, Adrian M. Gomez, Jeremy C. Borniger, Lucas Cheadle

## Abstract

To survive, organisms must adapt to a staggering diversity of environmental signals, ranging from sensory information to pathogenic infection, across the lifespan. At the same time, organisms intrinsically generate biological oscillations, such as circadian rhythms, without input from the environment. While the nervous system is well-suited to integrate extrinsic and intrinsic cues, how the brain balances these influences to shape biological function system-wide is not well understood at the molecular level. Here, we demonstrate that the cytokine receptor Fn14, previously identified as a mediator of sensory experience-dependent synaptic refinement during brain development, regulates neuronal activity and function in adult mice in a time-of-day-dependent manner. We show that a subset of excitatory pyramidal (PYR) neurons in the CA1 subregion of the hippocampus increase Fn14 expression when neuronal activity is heightened. Once expressed, Fn14 constrains the activity of these same PYR neurons, suggesting that Fn14 operates as a molecular brake on neuronal activity. Strikingly, differences in PYR neuron activity between mice lacking or expressing Fn14 were most robust at daily transitions between light and dark, and genetic ablation of Fn14 caused aberrations in circadian rhythms, sleep-wake states, and sensory-cued and spatial memory. At the cellular level, microglia contacted fewer, but larger, excitatory synapses in CA1 in the absence of Fn14, suggesting that these brain-resident immune cells may dampen neuronal activity by modifying synaptic inputs onto PYR neurons. Finally, mice lacking Fn14 exhibited heightened susceptibility to chemically induced seizures, implicating Fn14 in disorders characterized by hyperexcitation, such as epilepsy. Altogether, these findings reveal that cytokine receptors that mediates inflammation in the periphery, such as Fn14, can also play major roles in healthy neurological function in the adult brain downstream of both extrinsic and intrinsic cues.

**Highlights:**

- Neuronal activity induces *Fn14* expression in pyramidal neurons of the hippocampus
- Fn14 constrains neuronal activity near daily transitions between light and dark
- Loss of Fn14 lengthens the endogenous circadian period and disrupts sleep-wake states and memory
- Microglia contact excitatory synapses in an Fn14-dependent manner

## Introduction

Despite the long-held view of the nervous system as an immunologically privileged site, interactions between immune cells and neurons via cytokine signaling are now known to be integral to neural circuit development in the early postnatal brain^1^. For example, emerging work suggests that brain-resident immune cells, microglia, not only protect the brain from injury and disease, but also influence its development under normal physiological conditions^2,3^. While microglia contribute to multiple developmental processes, their most well-defined role is to remove excess or developmentally transient synapses via phagocytic engulfment or through the directed release of secreted factors onto neurons^4–7^. The competitive removal of a subset of immature synapses by microglia facilitates the strengthening and maintenance of a separate cohort of synapses, thereby driving circuit maturation. Furthermore, neurons themselves express numerous cytokines, cytokine receptors, and other immune-related signaling proteins, including Major Histocompatibility Complex (MHC) class I molecules and components of the classical complement cascade, which localize to developing synapses to mediate their elimination, remodeling, or strengthening via both microglia-dependent and microglia-independent mechanisms^8–10^. Thus, cytokines and their receptors are essential for brain development.

While the removal of excess synapses via cytokine signaling between microglia and neurons is critical for brain development, this process can become inappropriately heightened during aging, leading to the removal of mature synapses and eliciting cognitive decline in neurodegenerative conditions such as Alzheimer’s disease (AD)^11–13^. *A key conceptual link that is missing is an understanding of how cytokines operate within the mature brain to mediate its function and plasticity in the absence of disease*. To this point, several observations suggest that these pathways may be uniquely poised to play important roles in the adult brain. For example, like the immune system, the brain is a heterogeneous tapestry of diverse cell types which communicate with one another across spatial and temporal scales. Cytokine signaling molecules represent a promising mechanism to mediate interactions between brain cells that are not in direct contact, in part because these factors can be expressed in direct response to environmental cues^14,15^. In addition, just as synapses in the developing brain undergo dynamic changes in number, structure, and physiology, synapses are similarly remodeled in the mature brain to mediate the adaptation of neural circuits to dynamic changes in the environment^16,17^. Thus, the same immune-related mechanisms that regulate synaptic remodeling during development may also regulate this process in adulthood. Finally, there are likely to be evolutionarily conserved benefits of the immune system and the nervous system sharing a molecular language in the form of cytokines and their receptors, such as the facilitation of interactions between the body and the brain. However, whereas the neuronal populations that integrate and encode inflammatory signals in the periphery are beginning to be identified^18,19^, the molecular motifs that mediate this integration are not known. Thus, determining whether cytokines orchestrate mature brain function, and the specific ways in which they do so, is an important next step in elucidating the nature and importance of neuro-immune communication within the brain and beyond.

Among cytokine pathways that may be ideally poised to mediate adult brain function, the TWEAK-Fn14 pathway has emerged as a promising candidate. In this pathway, the Tumor necrosis factor (TNF) family cytokine TWEAK (TNF-associated weak inducer of apoptosis) binds to the TNF receptor family member Fn14 (Fibroblast growth factor inducible protein 14 kDa), thereby eliciting local cellular remodeling events alongside changes in gene expression that underlie processes such as inflammation, tissue regeneration, and angiogenesis^20–24^. Although Fn14 expression was previously thought to be low in the healthy brain, we recently identified a requirement of TWEAK-Fn14 signaling for the refinement of visual circuit connectivity between the retina and the dorsal lateral geniculate nucleus (dLGN) of the thalamus^7,25,26^. Homing in on a critical period of sensory experience-dependent plasticity that takes place during the third week of life, we found that, in response to acute visual stimulation, Fn14 is expressed at synapses between retinal ganglion cells (RGCs) and thalamic neurons of the dLGN^27,28^. When microglia release TWEAK onto synapses containing Fn14, these synapses are structurally disassembled and eliminated, allowing the synapses that are not exposed to soluble TWEAK to mature appropriately^7,25^. Thus, Fn14 acts as a sensor of visual information during circuit development, thereby mediating the impact of environmental cues on the connectivity of the brain. However, whether and how Fn14 mediates mature brain function was not known.

In this study, we harnessed the TWEAK-Fn14 pathway as a molecular handle to shed light on the roles of cytokine signaling in the mature brain. We found that Fn14 expression is dynamically upregulated in a subset of glutamatergic pyramidal (PYR) neurons in the CA1 subregion of the hippocampus, an area that mediates learning and memory, in response to neuronal activity. Upon its expression in active neurons, Fn14 functions to restrict their excitability, likely returning the circuit to a homeostatic state. Remarkably, the modulation of neuronal activity by Fn14 is most prominent near daily transitions between light and dark, suggesting the possibility of a circadian component to Fn14 function. Indeed, behavioral and neurophysiological analyses uncovered a role for Fn14 in sensory-cued and spatiotemporal memory, sleep-wake balance, and circadian rhythms *in vivo*. These data reveal an essential role for Fn14 in mature brain function, indicating that cytokine receptors that mediate inflammation in the periphery can also orchestrate core neurobiological processes that impact organismal health and survival as a whole. In combination with the known roles of TWEAK and Fn14 in sensory-dependent phases of brain development, these data suggest that Fn14 is poised to integrate the effects of extrinsic and intrinsic stimuli in the mature brain.

## Results

### Excitatory glutamatergic neurons in the adult mouse brain express Fn14

To characterize the potential roles of Fn14 in adult brain function, we first asked whether Fn14 is expressed in the adult brain and, if so, which regions and cell types express it. Toward this end, we quantified *Fn14* mRNA expression in sagittal sections of the mouse brain at postnatal day (P)28, when brain maturation is nearing completion, and in the fully mature brain at P90 using single-molecule fluorescence *in <u>s</u>itu* <u>h</u>ybridization (smFISH, RNAscope). At both ages, we observed *Fn14* expression in a subset of cells across a diversity of brain structures. *Fn14* expression generally increased along an anterior-to-posterior axis, and was particularly high in the cerebellum where it was largely restricted to the granule cell layer. *Fn14* was also observed in the brain stem, the dLGN and other thalamic nuclei, and select cells in the hippocampus and cortex (Fig. 1A,B).

**Figure 1.**
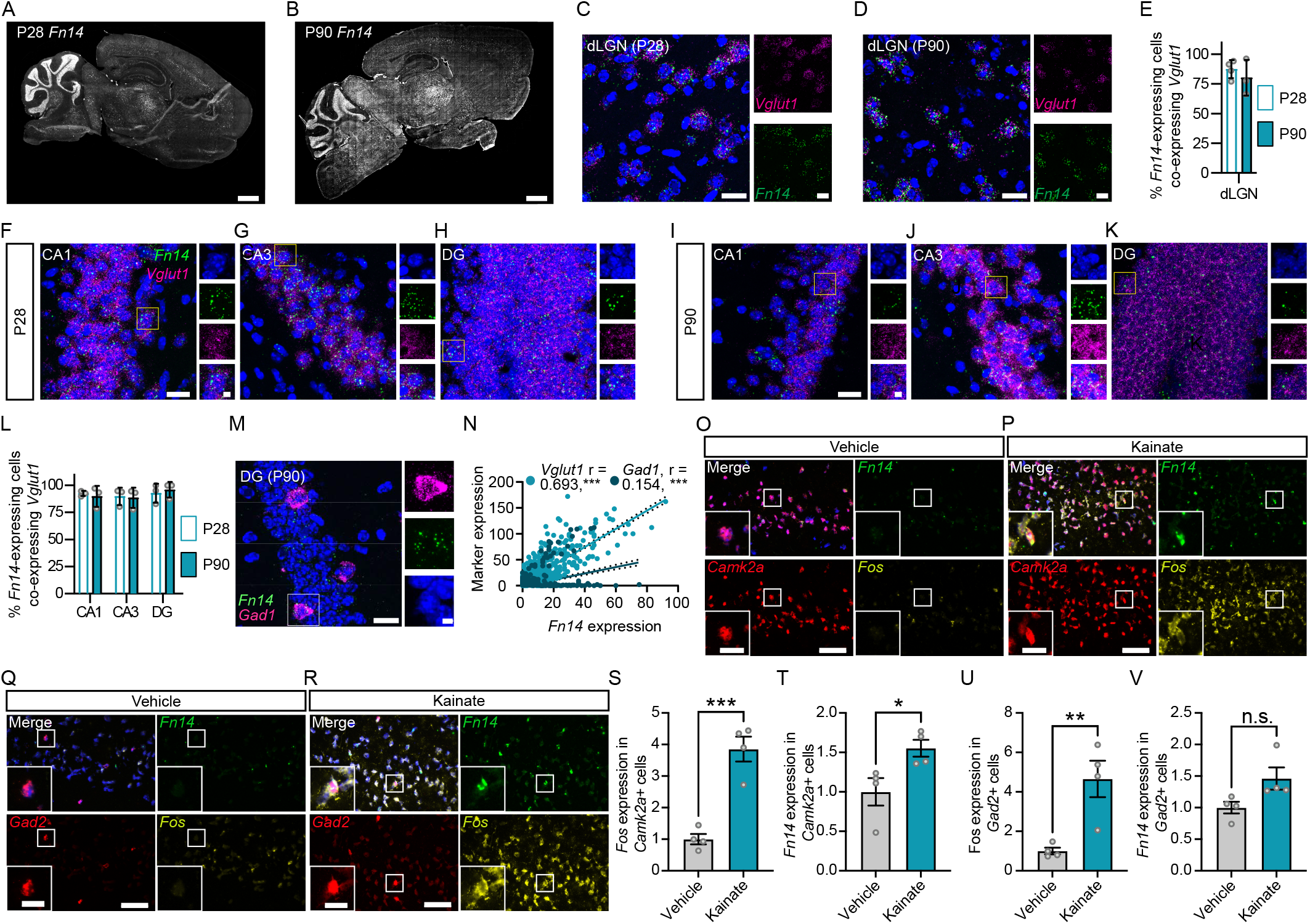
Neuronal activity induces *Fn14* expression in pyramidal neurons of hippocampal CA1. (A),(B) Confocal images of sagittal sections of the mouse brain at P28 (A) and P90 (B) subjected to single-molecule fluorescence *in situ* hybridization (smFISH) to label *Fn14* mRNA (white). Scale bars, 1 mm. (C),(D) High resolution confocal images of the dLGN in coronal sections from a P28 (C) and a P90 (D) mouse brain probed for *Fn14* (green) and the glutamatergic neuron marker *Vglut1* (magenta). DAPI shown in blue. Scale bars, 20 μm. (E) Quantification of the percentage of *Fn14*-expressing cells that also express *Vglut1* in the dLGN at P28 and P90 (unpaired Student’s T-test, p > 0.05). (F)-(H) Confocal images of CA1 (F), CA3 (G), and dentate gyrus ([DG]; H) subregions of the hippocampus in a coronal section from a P28 mouse brain probed for *Fn14* (green) and *Vglut1* (magenta). DAPI shown in blue. Scale bar, 20 μm. Inset scale bar, 5 μm. (I)-(K) Confocal images of CA1 (I), CA3 (J), and DG (K) regions of the hippocampus in a coronal section from a P90 mouse brain probed for *Fn14* (green) and *Vglut1* (magenta). DAPI shown in blue. Scale bar, 20 μm. Inset scale bar, 5 μm. (L) Quantification of *Fn14*-expressing cells that are positive for *Vglut1* in the hippocampus at both ages (Two-way ANOVA: region: p > 0.05, age: p > 0.05, interaction: p > 0.05). (M) Confocal image of *Fn14* (green) and the inhibitory neuron marker *Gad1* (magenta) in the DG at P90. Scale bar, 20 μm. Inset scale bar, 5 μm. (N) Scatter plot demonstrating the correlation between *Fn14* expression (x axis) and the expression of excitatory (*Vglut1*) or inhibitory (*Gad1*) neuron markers (y axis) in the hippocampus. Linear regression with slope comparison (***p < 0.001). Note the bimodal distribution for *Gad1* cells, suggesting that a defined subpopulation of inhibitory neurons may express Fn14. (O),(P) Confocal images of CA1 following smFISH for *Fn14* (green), the PYR neuron marker *Camk2a* (red), and the activity-dependent gene *Fos* (yellow) in mice exposed to vehicle (O) or kainate (P). Scale bar, 50 μm. Inset scale bar, 16 μm. (Q),(R) Confocal images of CA1 following smFISH for *Fn14* (green), the interneuron marker *Gad2* (red), and the activity-dependent gene *Fos* (yellow) in mice exposed to vehicle (Q) or kainate (R). Scale bar, 50 μm. Inset scale bar, 16 μm. (S),(T) Quantification of *Fos* (S) or *Fn14* (T) expression in *Camk2a*+ neurons in response to vehicle or kainate exposure, values normalized to vehicle. (U),(V) Quantification of *Fos* (U) or *Fn14* (V) expression in *Gad2*+ interneurons in response to vehicle or kainate exposure, values normalized to vehicle. Statistics for (S) – (V): Unpaired Student’s T-tests, ***p < 0.001; **p < 0.01; *p < 0.05; n.s. p > 0.05.

To identify the cell types that express *Fn14*, we assessed the colocalization of *Fn14* with the excitatory glutamatergic neuron marker *Vglut1* and the inhibitory neuron marker *Gad1* in two brain regions: the dLGN and the hippocampus. In the dLGN, a region in which *Fn14* expression is relatively high and in which we previously found Fn14 to mediate synaptic refinement^7,25^, the majority of *Fn14*+ cells (∼90%) at both P28 and P90 also expressed *Vglut1*, indicating that Fn14 is most highly expressed in excitatory neurons in this region (Fig. 1C-E). Next, we more closely examined *Fn14* expression in the hippocampus for the following reasons: (1) The hippocampus is essential for a plethora of critical brain functions that require synaptic plasticity, most notably learning and memory; (2) Hippocampal organization and connectivity have been well-characterized; and (3) Numerous physiological and behavioral paradigms have been developed to interrogate hippocampal circuitry and function. Quantification of *Fn14* expression in the three main interconnected hippocampal subregions (the dentate gyrus [DG], CA1, and CA3) at P28 and P90 revealed that, as in the dLGN, about 90% of *Fn14*+ cells also expressed the excitatory neuron marker *Vglut1*, which in the hippocampus labels pyramidal (PYR) neurons (Fig. 1F-L). Although *Fn14* expression was most frequently observed in excitatory neurons, we found *Fn14* in a subset of *Gad1*+ inhibitory neurons as well (Fig. 1M). Consistent with these observations, *Fn14* expression in the hippocampus was positively correlated with the expression of both *Vglut1* (r^2^ = 0.693; p < 0.001) and *Gad1* (r^2^ = 0.154; p < 0.001; Fig. 1N). Together, these results demonstrate that Fn14 is expressed by both excitatory and inhibitory neurons in the hippocampus, but that the majority of its expression is localized to excitatory cells. These data raised the possibility that Fn14 mediates hippocampal connectivity and function in the mature brain, possibly by operating within PYR neurons.

### Neuronal activity induces Fn14 expression in a subset of pyramidal neurons in hippocampal CA1

Memory encoding and retrieval are core functions of the hippocampus and occur, in part, through the coordination of activity-dependent gene programs that are induced in neurons downstream of synaptic activity. These gene programs encode molecules with direct roles in synaptic organization and remodeling, such as Fn14. Given its expression in excitatory and inhibitory neurons in CA1, along with prior evidence that Fn14 is upregulated in response to visual stimulation in the dLGN during development^25^, we hypothesized that Fn14 may be one of the activity-regulated molecules that facilitates the formation and storage of memory. If so, then the expression of Fn14 would be expected to be higher in active PYR neurons than in inactive neurons. To interrogate this possibility, we performed smFISH on the CA1 regions of the hippocampi of mice that had been systemically exposed to kainate ([10 mg/kg]; intraperitoneally; or water as vehicle control) for two hours. Kainate is a soluble compound that can cross the blood-brain barrier and bind a subset of glutamate receptors to induce the robust activation of neurons. In hippocampal slices from kainate-or vehicle-treated mice, we probed for *Fn14* along with the excitatory PYR neuron marker *Camk2a*, the inhibitory neuron marker *Gad2*, and *Fos*, an activity-regulated gene that served as a positive control^29^. As expected, *Fos* was significantly upregulated in both PYR and inhibitory neurons in CA1 following kainate exposure, validating kainate as a robust driver of neuronal activity-dependent transcription *in vivo* (Fig. 1O-S,U).

Similar to *Fos*, *Fn14* expression was also significantly higher in *Camk2a*+ excitatory neurons in kainate-treated mice than in vehicle-treated controls (Fig. 1O,P,T). Conversely, Fn14 expression within *Gad2*+ inhibitory neurons was not significantly altered by neuronal activation (Fig. 1Q,R,V and Fig. S1A). Two possible scenarios could give rise to the increase in *Fn14* expression observed in CA1 following kainate exposure: (1) the number of PYR neurons expressing *Fn14* could increase, or (2) the number of PYR neurons expressing *Fn14* may remain the same, but these neurons may express a greater amount of *Fn14* when activity is heightened. Our data revealed that the number of PYR neurons expressing *Fn14* was not altered by kainate exposure, supporting the latter interpretation that a subset of PYR neurons express *Fn14* more highly in response to activity (Fig. S1B). While kainate is a powerful stimulant that can activate neurons to an extent that is greater than what typically occurs *in vivo*^30^, we found that Fn14 expression was significantly higher in PYR neurons that expressed *Fos* (i.e. neurons that were recently activated) than in neurons that were Fos-negative, regardless of whether a mouse was exposed to kainate or vehicle (Fig. S1C). These observations are consistent with a scenario in which *Fn14* is transcribed in a distinct cohort of activated PYR neurons at a given time, potentially to mediate the encoding of memory in response to environmental cues that selectively activate this subset of neurons.

To validate these results, we assessed two whole-transcriptome datasets describing the transcriptional responses of hippocampal neurons to kainate *in vivo*^31,32^. In both datasets, *Fn14* was identified as being significantly induced by neuronal activation, confirming our findings (Fig. S1D-F). Interestingly, among the Tumor necrosis factor receptor (TNFR) superfamily members included in the study from Pollina *et al*, nine of the 18 genes encoding TNFRs exhibited either a significant upregulation (6 genes) or downregulation (3 genes) following kainate exposure compared to vehicle-treated controls (Fig. S1D). Thus, TNFRs other than Fn14 may also play important roles in the mature brain that have yet to be dissected. That said, among the six TNFRs that were upregulated by activity in the dataset, *Fn14* was by far the most strongly induced, underscoring that the functions of Fn14 in the brain are not likely to be redundant with the roles of other TNFRs.

### Fn14 is dispensable for learning but required for cued and spatial memory

Given the expression of *Fn14* in the hippocampus under normal physiological conditions and its induction in CA1 PYR neurons in response to neuronal activity, we next sought to determine whether Fn14 mediates hippocampal function. To this end, we analyzed learning and memory in a validated global Fn14 KO mouse line^25,33^ alongside WT littermates using two behavioral paradigms: cued fear conditioning (CFC) and Morris water maze (MWM). In the CFC task, we examined the abilities of Fn14 KO and WT mice to associate both an auditory (i.e. sensory) cue and a defined spatial context with a paired aversive foot shock (Fig. 2A). During the initial conditioning phase, when the foot shock was accompanied by an audible tone (75 dB; 2000 Hz) and a novel arena (striped walls and floor grating), both Fn14 KO and WT mice exhibited a stereotyped freezing response reflecting fear of the shock. Similarly, when mice of both genotypes were placed into a novel, unfamiliar context (a round arena with polka dotted walls) without a tone, they exhibited low levels of freezing. Next, the mice were subjected to probe trials in which they were exposed to (1) the shock-associated spatial context or (2) the shock-associated auditory tone in the absence of an accompanying foot shock. While Fn14 KO mice froze to a similar (though slightly lower) extent as WTs when re-exposed to the spatial context, they exhibited significantly less freezing when re-exposed to the auditory tone while in a novel environment (45.8 vs 66.9 seconds; Fig. 2B). This deficit could reflect an inability of mice lacking Fn14 to generalize their association of the tone with the foot shock to a new spatial context. Together, these data indicate that Fn14 likely contributes to the encoding and/or retrieval of memories, with the strongest deficits in Fn14 KO mice revolving around an inability to pair a sensory cue with an aversive stimulus.

**Figure 2.**
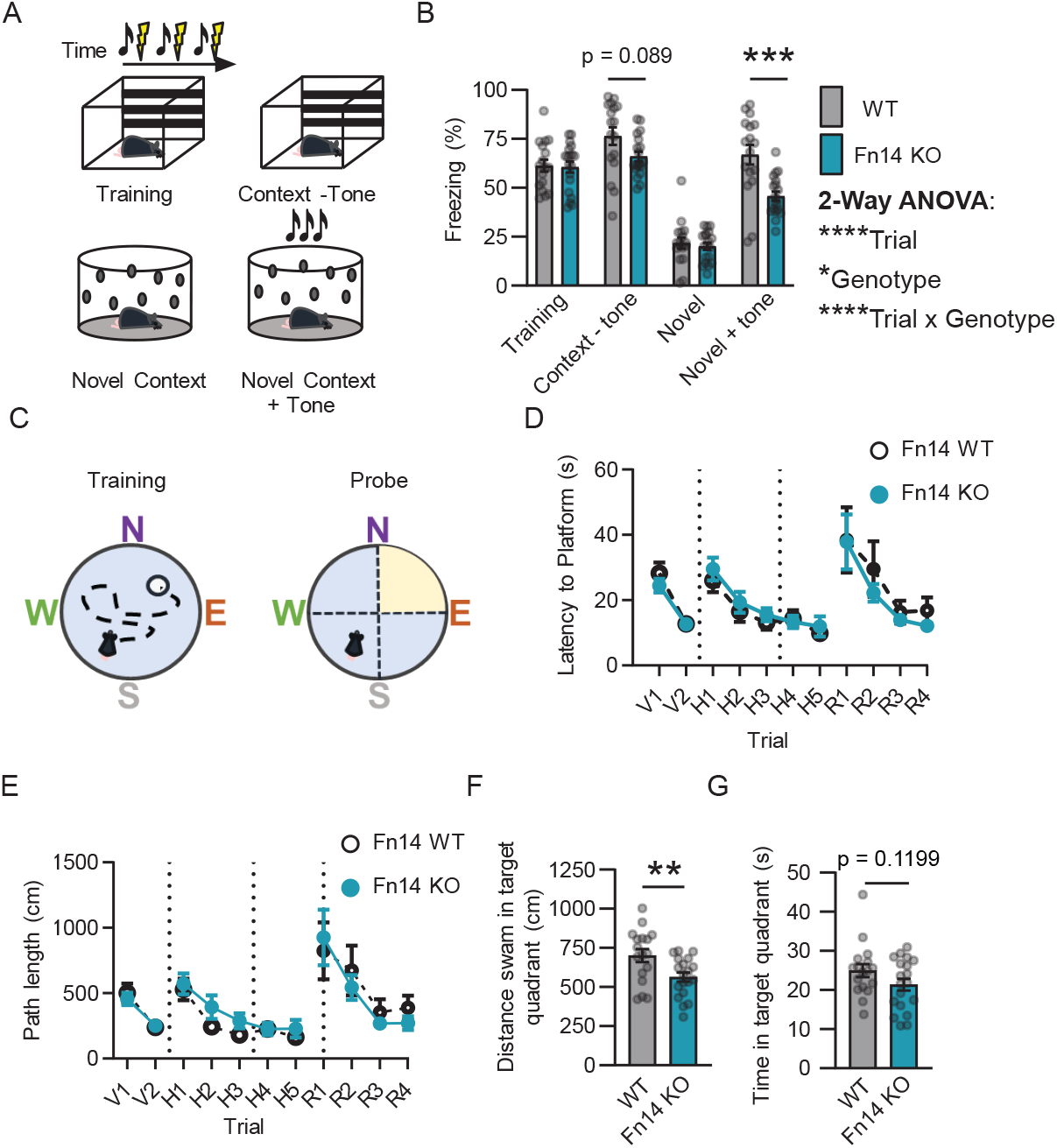
Fn14 is dispensable for learning but required for cued and spatial memory. (A) Diagram of the cued fear conditioning (CFC) paradigm. An auditory tone and a unique spatial context were initially paired with an aversive foot shock. The ability of mice to remember this association was later tested by exposing the mice to either the spatial context or the auditory tone in the absence of the shock. Freezing behavior, which mice exhibit when afraid, serves as a read-out for how well the mice remember the association between the context or tone and the shock. (B) Quantification of the percentage of time that mice spent freezing across all conditions (repeated measures ANOVA, trial: ****p < 0.0001; genotype: *p < 0.05; subject, trial x genotype: ****p < 0.0001). Bonferroni corrected multiple comparisons WT versus KO for Context (-) tone: p = 0.089; Novel context + tone: p < 0.001. (C) Diagram of Morris water maze (MWM) training and probe trials. (D) Latency to goal platform swam during MWM trials (repeated measures ANOVA with Šídák’s multiple comparisons test. Platform is <u>V</u>isible (V): genotype: p > 0.05, trial: p < 0.0001, trial x genotype: p > 0.05; platform is <u>H</u>idden (H): genotype: p > 0.05, trial: p < 0.0001, trial x genotype: p > 0.05; platform goal zone is <u>R</u>eversed (R): genotype: p > 0.05, trial: p < 0.001, trial x genotype: p > 0.05. (E) Path length swam by WT and Fn14 KO mice during the MWM test (repeated measures ANOVA with Šídák’s multiple comparisons test. V: genotype: p > 0.05, trial: p < 0.0001, trial x genotype: p > 0.05; H: genotype: p > 0.05, trial: p < 0.0001, trial genotype: p > 0.05; R: genotype: p > 0.05, trial: p < 0.001, trial x genotype: p > 0.05). (F) Distance swam by mice in the target quadrant during the probe trial (cm; unpaired Student’s T-Test, **p < 0.01). (G) Time spent in target quadrant during probe trial (s; unpaired Student’s T-Test, p > 0.05). For all analyses: n = 17 WT and 19 Fn14 KO mice. **p < 0.01, ***p < 0.001.

Because impairments in the CFC task could reflect functional changes in the amygdala or the frontal cortex in addition to the hippocampus, we next examined whether the loss of Fn14 would have a similar effect on a more purely hippocampal-dependent spatial learning task, the MWM. In this task, the mice were placed in a round pool with each cardinal direction marked by a distinctive shape and color to allow for spatial mapping of the arena (Fig. 2C). During the initial training stage, WT and Fn14 KO mice were both able to effectively locate a visible goal platform. After mice were trained to perform the task, the water in the pool was made opaque and the goal platform submerged, promoting the use of spatial orientation-based strategies for locating the goal platform, rather than the platform itself^34^. In all trials in which the platform was hidden, WT and Fn14 KO mice learned to find the platform equally well as revealed by their similar latencies to reach the platform and the lengths of the paths that they took to reach it (Fig. 2D,E). Thus, as also demonstrated by the results of the CFC task, loss of Fn14 does not have a strong observable effect on learning.

To specifically assess spatial memory function, we next tested whether, after a period of 24 hours, the mice remembered the location of the hidden platform. When the platform was removed from the pool in probe trials, WT mice swam a significantly greater distance in the quadrant where the platform was previously hidden than Fn14 KO mice by about 25%, suggesting that WT mice were able to remember the location of the platform while mice lacking Fn14 did so less effectively (Fig. 2F). As expected, the decreased distance swam in the goal quadrant by the Fn14 KO mice corresponded with a trend toward less time spent in the target quadrant (Fig. 2G). These deficits were not caused by an impairment in visual or motor function, as WT and Fn14 KO mice swam an equal distance overall during the probe trial, and Fn14 KO mice exhibited normal visual acuity as assessed by optomotor testing (Fig. S2). Following the probe trials, the goal platform was reintroduced into the pool, but now in the opposite quadrant of the arena. Just as in the hidden trials, both WT and Fn14 KO mice were able to learn the new reversed goal zone equally well, again suggesting that Fn14 does not affect the acquisition of new information (Fig. 2D,E). Thus, Fn14 is dispensable for learning but required for memory.

### Fn14 dampens PYR neuron activity in vivo

The observation that PYR neurons in hippocampal CA1 induce *Fn14* expression in response to neuronal activity, along with the memory deficits observed in Fn14 KO mice, suggests that CA1 may be a locus of Fn14 function in the mature brain. Thus, we next sought to examine the effects of genetic ablation of Fn14 on the activity states and physiological properties of CA1 PYR neurons in awake, behaving mice using fiber photometry. Briefly, this approach employs the viral transduction of excitatory PYR neurons in CA1 with the genetically encoded calcium indicator GCaMP6f downstream of the Camk2a promoter, allowing for specific transduction of excitatory CA1 PYR neurons^35^. Thus, expression of GCaMP6f is restricted to PYR neurons genetically via the promoter and spatially due to the precise stereotaxic injection of the AAV-Camk2a-GCaMP6f virus into CA1. An optic fiber is then implanted over CA1 to detect changes in the amount of internal Ca^2+^ via changes in GCaMP6f fluorescence (ΔF/F) which serves as a surrogate read-out of aggregate neuronal activity. Ca^2+^ transients or events, defined as temporal loci at which changes in ΔF/F meet a minimum threshold, are then quantified as a reflection of the activity states of the cells being recorded.

To determine whether Fn14 influences the activity of PYR neurons *in vivo*, we assessed the maximum amplitudes, as well as the frequency, of Ca^2+^ transients in CA1 PYR neurons from Fn14 KO and WT littermates over a 24-hour period during normal home cage behavior (Fig. 3A,B). While we did not observe differences in the maximum amplitude (ΔF/F) of Ca^2+^ events between genotypes (Fig. 3C,D), Fn14 KO mice exhibited a higher frequency of Ca^2+^ transients than WT controls (Fig. 3E,F). This result suggests that PYR neurons (or, most likely, a subset thereof) are more active in Fn14 KO mice than in WT littermates. Next, we assessed how differences in PYR neuron activity between WT and Fn14 KO mice fluctuated across 24 hours. Strikingly, we found that the extent to which the loss of Fn14 increased Ca^2+^ transient frequency varied substantially by time-of-day. For example, CA1 PYR neurons in the KO demonstrated the most robust increase in Ca^2+^ event frequency over PYR neurons in WT mice at Zeitgeber time (ZT) 11, an hour before lights are turned off in the mouse facility and mice generally transition from less active to more active states (Fig. 3A,E). Furthermore, when we isolated and aggregated the frequency of Ca^2+^ events exhibited during the light and dark periods (light = ZT 0-11; dark = ZT 12-23), we found that Fn14 KO mice exhibited an increase in Ca^2+^ transient frequency only during the dark phase, when mice are more active (Fig. 3F). These data suggest that Fn14 constrains the activity of PYR neurons under normal physiological conditions in a time-of-day-dependent manner.

**Figure 3.**
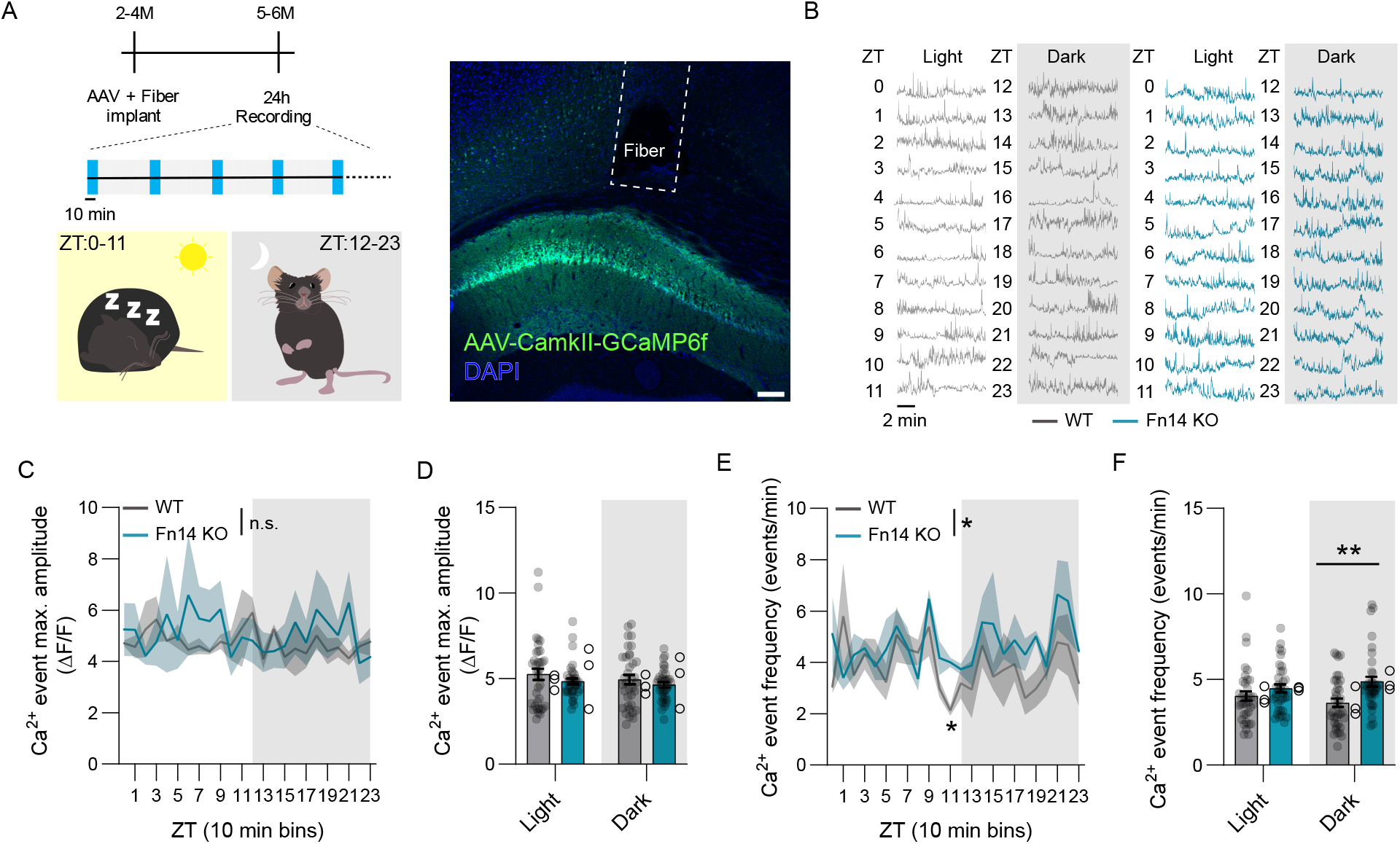
Fn14 dampens pyramidal neuron activity in a time-of-day-dependent manner. (A) Schematic of the experimental timeline with an example confocal image of GCaMP6f expression in CA1 and the optic fiber tract right above CA1. Scale bar, 100 µm. ZT, Zeitgeber time (mouse’s subjective time-of-day). Blue bars, 10-minute recording periods. M, months of age. H, hours. (B) Example 10-minute binned calcium traces (ΔF/F) from a representative WT and Fn14 KO mouse, recorded every hour (ZT0-23) over a single day. (C) Maximum ΔF/F signal over each time bin (repeated measures 2-way ANOVA: Time: p > 0.05, genotype: p > 0.05, interaction: p > 0.05). (D) Analysis of maximum ΔF/F signal during the light and dark phases of the day plotted as z scores (repeated measures 2-way ANOVA: Time: p > 0.05, genotype: p > 0.05, interaction: p > 0.05). (E) Ca^2+^ event frequency in WT and Fn14 KO mice over a 24-hour recording period (repeated measures 2-way ANOVA: Time: p > 0.05, genotype: p < 0.05, interaction p > 0.05, with Tukey post-hoc test: *p < 0.05 at ZT11). (F) Quantification of the Ca^2+^ event frequency during the light (ZT0-11) and dark (ZT12-23) phases of the day (repeated measures ANOVA: Time: p > 0.05, genotype: p < 0.01, interaction: p > 0.05, with Tukey post-hoc test: **p < 0.01 during the dark phase). For all analyses, n = 36 traces from 3 mice per genotype. Line graphs and histograms show mean ± S.E.M. while histograms show both acquisitions (closed circles) and within mouse averages (open circles).

We next sought to corroborate the finding that neurons lacking Fn14 are more active than their WT counterparts at the molecular level by assessing the expression and activation of the activity-dependent transcription factors Fos and Jun in whole brain homogenates from Fn14 KO and WT mice using ELISAs. Fos and Jun are members of the AP1 family of transcription factors that are activated by neuronal excitation, and they are also targets of the MAPK and JNK/p38 pathways which can be directly regulated by TWEAK and Fn14^21,36^. Consistent with neurons being overly active in the absence of Fn14, we observed significantly increased levels of phosphorylated (i.e. more active) versus unphosphorylated (i.e. less active) Jun, and a trend toward increased levels of Fos protein, in the brains of Fn14 KO mice compared to WT (Fig. S3A,B). One possible interpretation of these data is that Fn14 constrains neuronal activity, at least in part, by modulating the activation of AP1-mediated transcription. Although the gene programs that may be activated downstream of Fn14-AP1 interactions in neurons are yet to be defined, a candidate-based approach revealed significantly decreased expression of *Scn1a* in the brains of Fn14 KO compared to WT mice (Fig. S3C). *Scn1a* encodes a sodium channel subunit that regulates neuronal excitability, and mutations in the human *SCN1A* gene are among the strongest genetic drivers of epilepsy and seizures^37^. Together, these data provide physiological and molecular evidence that Fn14 dampens the activity of hippocampal neurons in the brain, possibly through a transcriptional mechanism that mirrors how Fn14 regulates inflammation in peripheral cells.

### Fn14 restricts the length of the endogenous circadian period and influences sleep-wake states

While mice lacking Fn14 exhibited increased PYR neuron activity over WT mice on average, when we plotted the fiber photometry data across the 24-hour recording period, we noted that the most striking difference between Fn14 KO and WT mice occurred an hour before the daily transition from the light phase to the dark phase (Fig. 3E). This led us to consider the possibility that the functions of Fn14 in the adult brain may be related to circadian rhythms. Circadian rhythms are endogenously generated biological oscillations that are expressed in almost all taxa, and closely match a cycle period of 24 hours. Though intrinsically determined, circadian rhythms can be modulated by environmental cues such as light, which is important for allowing organisms to match internal states to changes in the environment. Given that Fn14 constrains activity in a time-of-day-dependent manner, we asked whether circadian rhythms were altered in mice due to loss of Fn14. To this end, we employed a locomotor wheel-running assay to map active and inactive states of WT and Fn14 KO mice either in a normal 12-hour/12-hour light/dark cycling environment (as is found in most standard animal facilities) or in complete darkness for 24 hours a day. Measuring wheel-running in mice is an established method of interrogating the behavioral output of the circadian clock. Since mice are nocturnal and run on the wheel mostly when awake, they exhibit distinct periods of running wheel activity in a standard environment that sync up with the dark phase of the light/dark cycle. On the other hand, removing light cues allows for the unveiling of the mouse’s endogenous period (i.e. the free-running period) as light information is no longer available to entrain circadian rhythms to cues in the external environment. By measuring the running wheel activity of Fn14 KO and WT mice under normal light/dark conditions, we found that both KOs and WTs maintained the expected 24-hour circadian period (Fig 4A (top),B and Fig. S4). However, when the activity of mice was measured in constant darkness, Fn14 KO mice maintained an endogenous activity period that was significantly longer than that of their WT littermates (Fig. 4A (bottom),C). Thus, Fn14 may play a role in confining the length of the endogenous circadian period in mice, suggesting a role for cytokine signaling in the orchestration of internally driven oscillations that are initiated in the brain.

**Figure 4.**
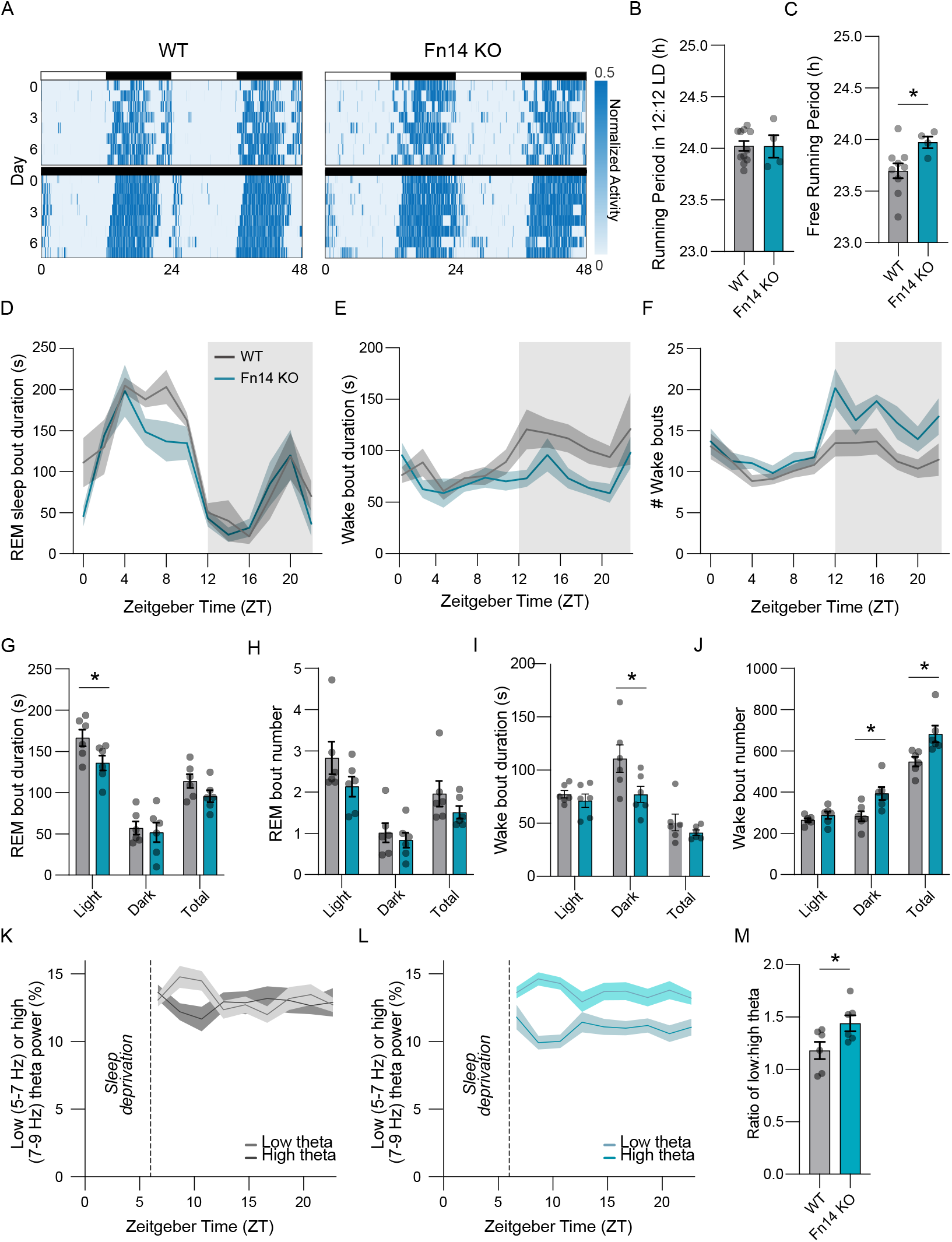
Fn14 regulates the length of the endogenous circadian period and modulates sleep-wake states in mice. (A) Representative actograms from WT and Fn14 KO mice under normal 12h:12h light/ dark conditions (top) as well as in constant darkness (bottom). Normalized running wheel activity is represented based upon the scale to the right, with higher levels of activity presenting as darker shades of blue. When housed in constant darkness, WT mice exhibit left-shifted activity periods reflective of a shorter circadian rhythm, whereas this left shift is absent in Fn14 KO mice. (B) Periodicity of running wheel activity under normal light/dark conditions (unpaired Student’s T-test: p > 0.05). (C) Free-running period during constant darkness, representative of the mouse’s innate circadian rhythm (unpaired Student’s T-test: p < 0.05). For (B) and (C), n = 11 WT and 4 Fn14 KO mice. (D-F) EEG/EMG analysis of REM sleep bout duration (D), number of REM sleep bouts (E), and number of wake bouts (F) for Fn14 KO and WT mice plotted over the 24-hour recording period. (G) Quantification of REM sleep bout duration (seconds) during the light phase, the dark phase, and over the full 24-hour period (total). (H) Quantification of REM coverage within both phases and over the full 24-hour period (total). (I) Mean number of REM bouts within both phases and over the full 24-hour period. (J) Mean number of wake bouts within both phases and over the full 24-hour period. (K) low (light gray) and high (dark gray) theta frequency bands following sleep deprivation in WT mice. (L) low (light teal) and high (dark teal) theta frequency bands following sleep deprivation in Fn14 KO mice. (M) Quantification of the ratio of low to high theta frequency in WT and Fn14 KO mice. Statistics for (G) – (J) and (M): multiple unpaired Student’s T-Tests, *p < 0.05.

Circadian rhythms play an important role in brain function and whole-body physiology, and are particularly critical for the regulation of oscillations in large-scale brain activity and sleep-wake states^38,39^. Thus, we next asked how the loss of Fn14 impacts sleep and wakefulness in mice by performing chronic, wireless electroencephalogram /electromyography (EMG/EEG) telemetry with concurrent activity monitoring during normal home cage behavior over a period of 48 hours. By correlating behavioral activity with EEG/EMG data using a standardized approach^40^, we quantified Non-Rapid Eye Movement (NREM) sleep characteristics, Rapid Eye Movement (REM) sleep characteristics, and wakefulness in Fn14 KO and WT mice (Fig. 4D-J). While the organization of NREM sleep was normal in Fn14 KO mice, the average duration of REM sleep bouts was decreased in the absence of Fn14 (Fig. 4D,G). We also observed a trend toward a decrease in the number of REM sleep bouts in Fn14 KO mice, suggesting that mice lacking Fn14 experience less REM sleep than their WT counterparts (Fig. 4H). Moreover, consistent with our finding that Fn14 constrains neuronal activity in a time-of-day-dependent manner, these decreases in REM sleep in Fn14 KO mice were restricted to the light cycle. We next evaluated the organization of waking behavior exhibited by Fn14 KO and WT mice across the recording period. We found that wake bout durations were lower in Fn14 KO mice than in WT mice during the dark phase, but that the number of wake bouts was simultaneously increased, potentially in an effort to compensate for the decreased bout duration (Fig. 4E,F,I,J). Alongside the decrease in REM sleep experienced by mice lacking Fn14, these changes in wake bout number and duration suggest that sleep-wake states in Fn14 KO mice are, at least to some extent, fragmented.

After recording sleep-wake states in mice under normal conditions, we applied a sleep deprivation protocol to determine whether Fn14 plays a role in the re-establishment of sleep-wake patterns following forced disturbances in sleep. Briefly, we subjected mice to ‘gentle handling’ for the first six hours of the light cycle, when mice spend most of their time sleeping. Recovery sleep and wake data were then recorded over the subsequent 18 hours (Fig. S5). Analyzing EEG/EMG data following an acute 6-hour sleep deprivation protocol, we found that Fn14 KO mice exhibited higher low-to-high theta band ratios during wakefulness than WT mice during the recovery period (Fig. 4K-M). As the prevalence of low theta (5-7 Hz) to high theta (7-9 Hz) activity during wakefulness is thought to be related to sleep propensity^41^, or the drive to attain sleep following a period of wakefulness, this result suggests that Fn14 KO mice were more tired, or fatigued, following sleep deprivation than their WT counterparts. This finding is consistent with the baseline fragmentation of sleep and the impairments in memory displayed by Fn14 KO mice (Fig. 2). Overall, these data provide evidence that Fn14 influences circadian rhythms and sleep/wake states *in vivo*.

### Microglia contact fewer, but larger, excitatory synapses in the absence of Fn14

A unique population of brain-resident immune cells, microglia, are the predominant expressers of the Fn14 ligand TWEAK in the brain^25,42^. In the developing visual system, microglia-derived TWEAK converges upon synaptic Fn14 to structurally disassemble a subset of synapses, thereby driving circuit maturation. Given the ability of Fn14 to constrain neuronal activity in CA1, we hypothesized that Fn14 may recruit microglia to remove, weaken, or otherwise modify synaptic inputs onto PYR cells, thereby dampening their activity. Consistent with this possibility, we found by immunofluorescence that microglia contact significantly fewer vGluT1+ synapses in hippocampal CA1 in Fn14 KO compared to WT mice (Fig. 5A-C). This result suggests that Fn14 may recruit microglia to disassemble excitatory synapses onto PYR cells, similar to the roles of this pathway in visual circuit development^7^. In line with this possibility, the vGluT1+ synapses that were contacted by microglia in the Fn14 KO mouse were significantly larger than those contacted by microglia in WT mice (Fig. 5D). One possible interpretation of this result is that the synapses that were contacted by microglia in the absence of Fn14 were less likely to be in a state of disassembly than the smaller synapses contacted by microglia in the WT. A similar analysis of contacts between microglia and vGat+ inhibitory synapses revealed that, while microglia contacted the same number of vGat+ synapses in KO and WT mice, the vGat synapses contacted by microglia were also larger in Fn14 KO mice than in WT littermates (Fig. 5E,F). These observations suggest that TWEAK-Fn14 signaling from microglia to neurons may modify synapses onto PYR neurons in CA1, possibly to facilitate the constraint of PYR neuron activity.

**Figure 5.**
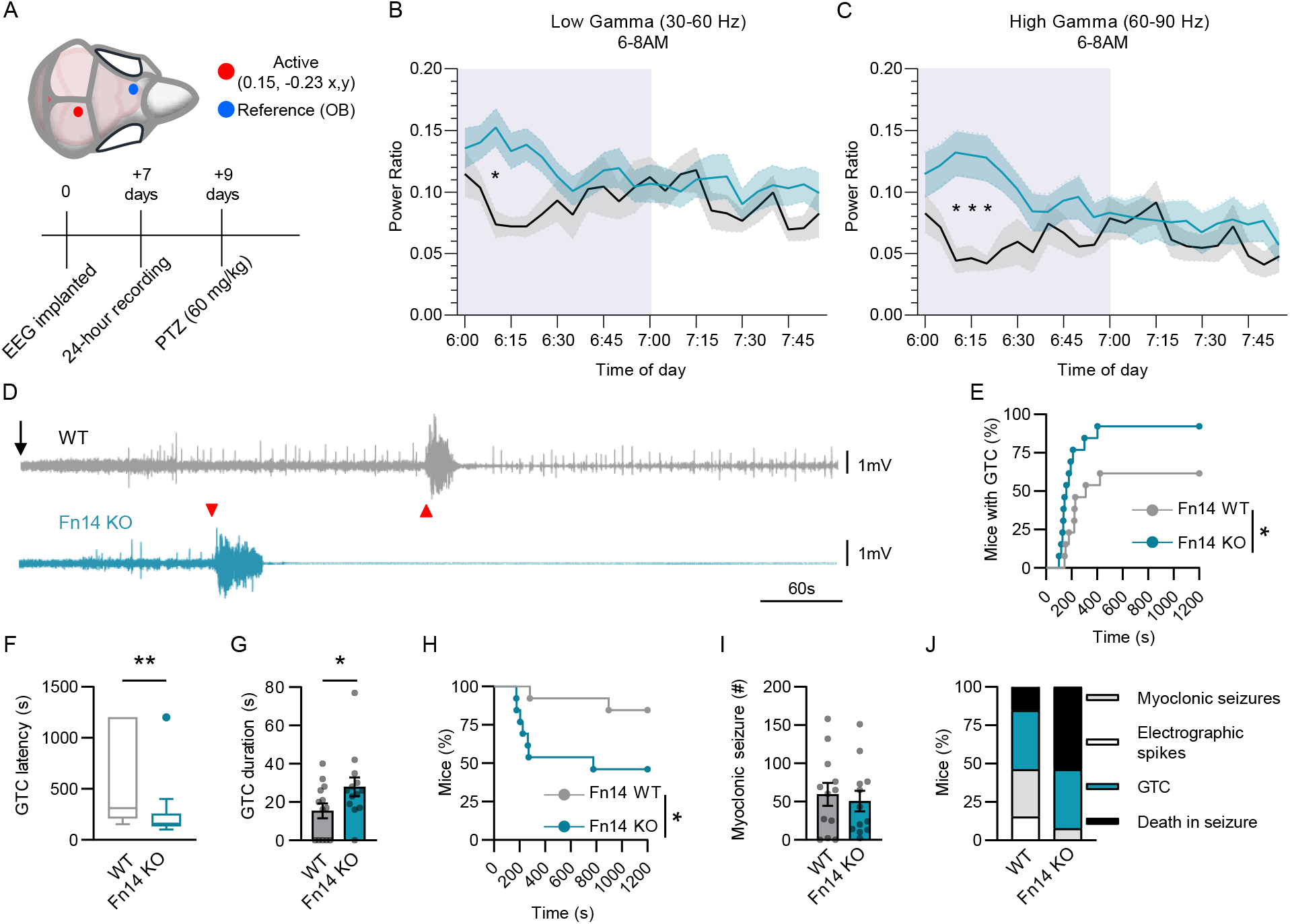
Microglia contact fewer but larger excitatory synapses in the absence of Fn14. (A),(B) Example reconstructions of microglia (Iba1, green) surrounded by excitatory synapses (Vglut1, magenta) and inhibitory synapses (Vgat, cyan) in hippocampal CA1. Microglia reconstructed from a WT (A) and an Fn14 KO mouse (B). Confocal images from which microglia and synaptic inputs were reconstructed are shown on the right. Scale bars, 5 μm. (C),(E) Quantification of the number of Vglut1+ excitatory synapses (C) or Vgat+ inhibitory synapses (E) contacted by microglia in Fn14 KO and WT mice. Log-scales were used because they best fit the distribution of the data. (D),(F) Quantification of the average volume of Vglut1+ synapses (D) or Vgat+ synapses (F) contacted by microglia. For (C)-(F), Mann-Whitney Tests, *p < 0.05, **p < 0.01, ***p < 0.001. Individual datapoints represent microglia while open circles indicate mouse averages; n = 45/50 microglia from 3 WT/4 KO mice.

### Loss of Fn14 increases seizure severity and seizure-related mortality

Neuronal activity promotes memory through the induction of plasticity mechanisms, i.e. long-term potentiation and/or long-term depression, at synapses. However, inappropriately prolonged activity can have deleterious effects that can lead to neuronal dysfunction and eventually cell death through excitotoxicity^43^. Therefore, the brain employs various mechanisms to gate increases in neuronal activity to balance excitation and inhibition and to maintain circuit homeostasis. The disruption of these activity-gating mechanisms has the potential to lead to the correlated hyperexcitability of neurons which, in extreme cases, causes seizure activity characteristic of epilepsy.

One proposed role for cytokine signaling in the brain, and for microglia in general^44,45^, is to provide negative feedback on runaway neuronal activity, thereby protecting neurons from hyperexcitability. Given evidence that Fn14 constrains the activity of neurons in the healthy brain, we next sought to determine whether Fn14 is sufficient to protect circuits from hyperexcitability in a pathological context. To test this hypothesis, we first asked whether genetic ablation of Fn14 impacts brain activity on a macroscopic level. To address this question *in vivo*, we implanted EEG probes into the dorsal skulls (near where the hippocampus is located) of Fn14 KO mice and WT littermates and quantified the effect of loss of Fn14 on brain activity over a 48-hour period (Fig. 6A). These experiments revealed no differences in EEG spectral power between Fn14 KO and WT mice in delta (1-4 Hz), theta (4-8 Hz), alpha (8-12 Hz), beta (12-30 Hz), low gamma (30-60 Hz) or high gamma (60-90 Hz) frequency bands averaged across the 48-hour recording period (Fig. S6). Although averaged EEG values were not different between Fn14 KO and WT mice, the temporal resolution of these experiments allowed us to look more closely at how brain activity changed across the 48-hour recording session. In doing so, we observed a significant increase in low gamma activity in Fn14 KO mice compared to WT at about 6:15 AM (45 minutes before lights turned on) and an increase in high gamma activity around the same time (Fig. 6B,C). These data suggest that Fn14 constrains brain activity in a time-of-day-dependent manner with the biggest changes in activity occurring in the dark phase near the daily transition to the light phase, consistent with the results of our fiber photometry analysis (Fig. 3E,F).

**Figure 6.**
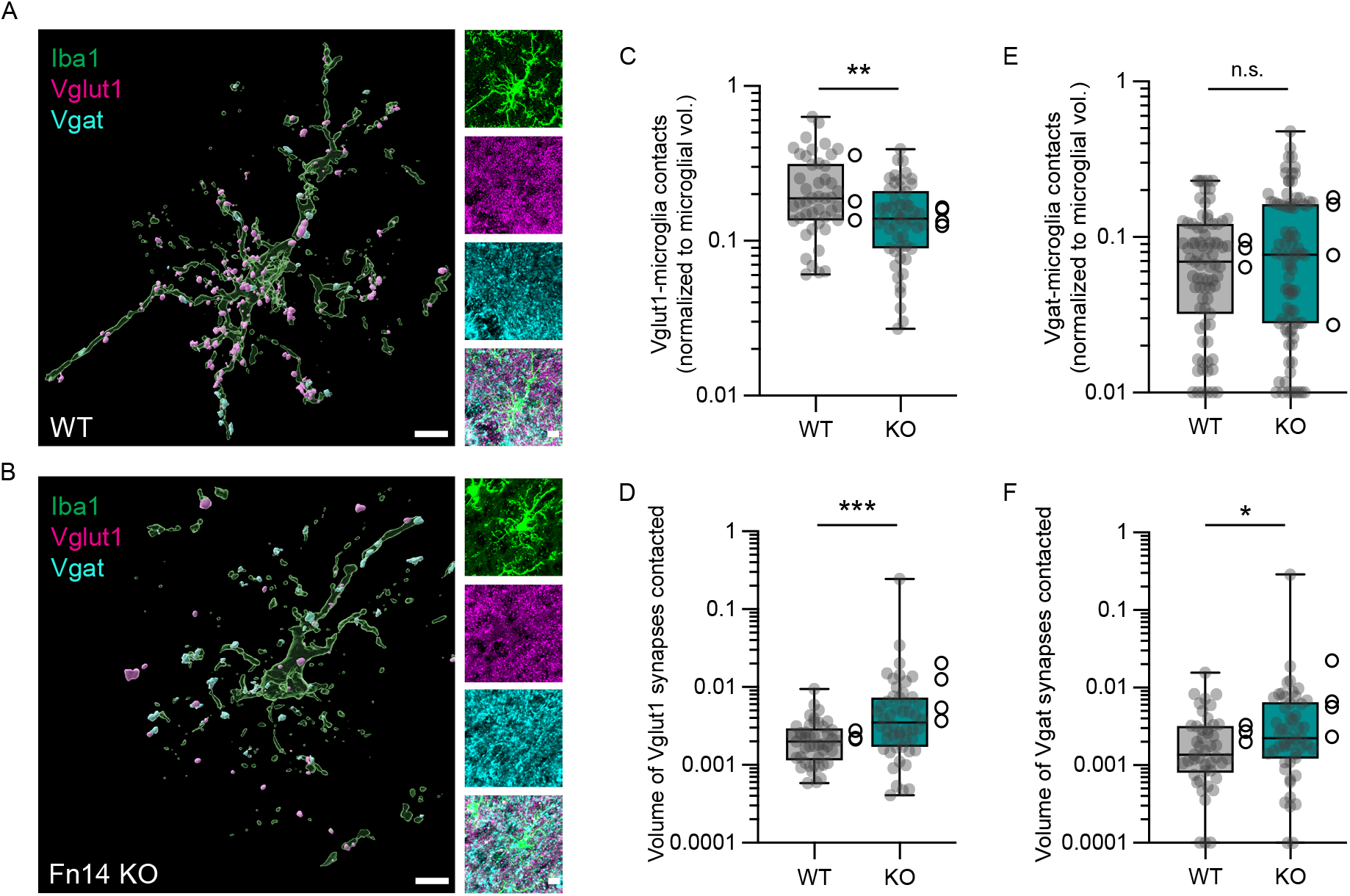
Fn14 is protective against chemically induced seizures. (A) Schematic of electroencephalogram (EEG) electrode placement and the experimental timeline. (B),(C) Traces (lines, mean; shaded areas, S.E.M.) of low gamma (B) and high gamma (C) activity between 6:00 and 8:00 AM. Lights on at 7:00 AM. (D) Example EEG traces from WT (gray) and Fn14 KO (teal) mice after PTZ injection (black arrow). Red triangles indicate the onset of general tonic clonic (GTC) seizures (WT: latency = 311 s, duration = 19.8 s; Fn14 KO: latency = 159 s, duration = 35 s). The Fn14 KO mouse died shortly after the GTC, demonstrated by the elimination of signal following the seizure. (E) Percentage of mice that had GTC seizures relative to the time course of the experiment (WT; n = 13 median = 311 s, Fn14 KO; n = 13, median = 159 s; Log-Rank test: *p < 0.05). (F) Latency between PTZ injection and GTC onset (Mann-Whitney test; **p < 0.01). (G) Duration of GTCs (unpaired Student’s T-test; *p < 0.05). (H) Mortality rate of Fn14 KO and WT mice following PTZ administration. Log-Rank test; *p < 0.05. (I) Number of PTZ-induced myoclonic seizures (Mann-Whitney test, p > 0.05). (J) The fraction of mice presenting with electrophysiological spikes (white), myoclonic seizures (grey), GTCs (teal), or death as their worst PTZ-induced outcome. Data presented as mean ± S.E.M. with data points representing individual mice or as the percentage of subjects, where applicable.

We next sought to determine whether, in the context of chemically induced seizures, Fn14 protects neurons from hyperexcitability by dampening their activity. To do so, we exposed Fn14 KO and WT mice to the GABA_a_ antagonist pentylenetetrazole (PTZ), a convulsant agent commonly used to elicit seizures through the dampening of inhibition onto excitatory hippocampal neurons which we have shown to inducibly express Fn14^46^. Intraperitoneal injection of PTZ (60 mg/kg) into Fn14 KO and WT mice time-locked with EEG recordings demonstrated profound differences in the responses of mice of each genotype to PTZ. Specifically, upon PTZ injection, Fn14 KO mice were more likely than WT littermates to develop general tonic clonic (GTC) seizures, and Fn14 KO mice developed GTCs at a shorter latency than WT mice (Fig. 6D-F). Furthermore, Fn14 KO mice exhibited a 152% increase in the duration of GTC seizures when compared to the GTCs measured in WT mice (Fig. 6G). Concurrent with the marked increase in GTC severity, Fn14 KO mice had a significantly higher mortality rate after PTZ challenge than WT mice, with about 50% of Fn14 KO mice dying as a result of seizure induction (Fig. 6H). There was no difference in the number of myoclonic seizures exhibited by Fn14 WT or KO mice, potentially due to the higher mortality rate of Fn14 KO mice (Fig. 6I). Lastly, loss of Fn14 led to a worse overall seizure phenotype as scored by a combination of recorded behavior, EEG activity, and mortality, suggesting that loss of Fn14 confers an increased susceptibility to acutely induced seizures that is extreme enough to cause death (Fig. 6J). Altogether, these functional data reveal that large-scale brain activity is heightened in the absence of Fn14 in a time-of-day-dependent manner, and that loss of Fn14 exacerbates seizure severity and worsens seizure outcomes following the acute dampening of inhibition. These results are consistent with a model in which Fn14 constitutes an activity-dependent feedback loop that protects neurons from hyperexcitability by dampening their activity.

## Discussion

In this study, we characterized the roles of the cytokine receptor Fn14 in mature brain function with a focus on the hippocampus, a structure that mediates learning and memory. We show that Fn14 is expressed in subsets of excitatory glutamatergic neurons throughout the brain including in the hippocampus, and that Fn14 expression is upregulated in a subset of PYR neurons in hippocampal CA1 in response to neuronal activity. In turn, Fn14 constrains the activity of neurons both under normal physiological conditions and in response to chemically induced seizures. These results suggest that Fn14 constitutes a molecular feedback mechanism that is turned on when a neuron becomes active then inhibits neuronal activity to return the neuron to a homeostatic state. Remarkably, Fn14 dampens neuronal activity most robustly near daily transitions between light and dark and during the dark phase, suggesting the possibility of a circadian component to Fn14 function. Indeed, mice lacking Fn14 exhibited significant aberrations in circadian rhythms and sleep-wake states, as well as deficits in cued and spatial memory (Fig. 7). Genetic ablation of Fn14 heightened the activation of AP1 transcription factors and decreased the expression of the epilepsy-related ion channel gene *Scn1a*, suggesting that Fn14 may mediate neuronal excitability at least in part through a transcriptional mechanism. On the other hand, microglia contacted fewer (but larger) excitatory synapses in CA1 in Fn14 KO compared to WT mice, indicating that Fn14 may recruit microglia to modify synapses acutely, thereby dampening PYR excitation. Altogether, these results reveal Fn14 as a coordinator of mature brain function, highlighting that molecules that mediate inflammation outside of the brain can contribute to sustaining neurological health across the lifespan.

**Figure 7.**
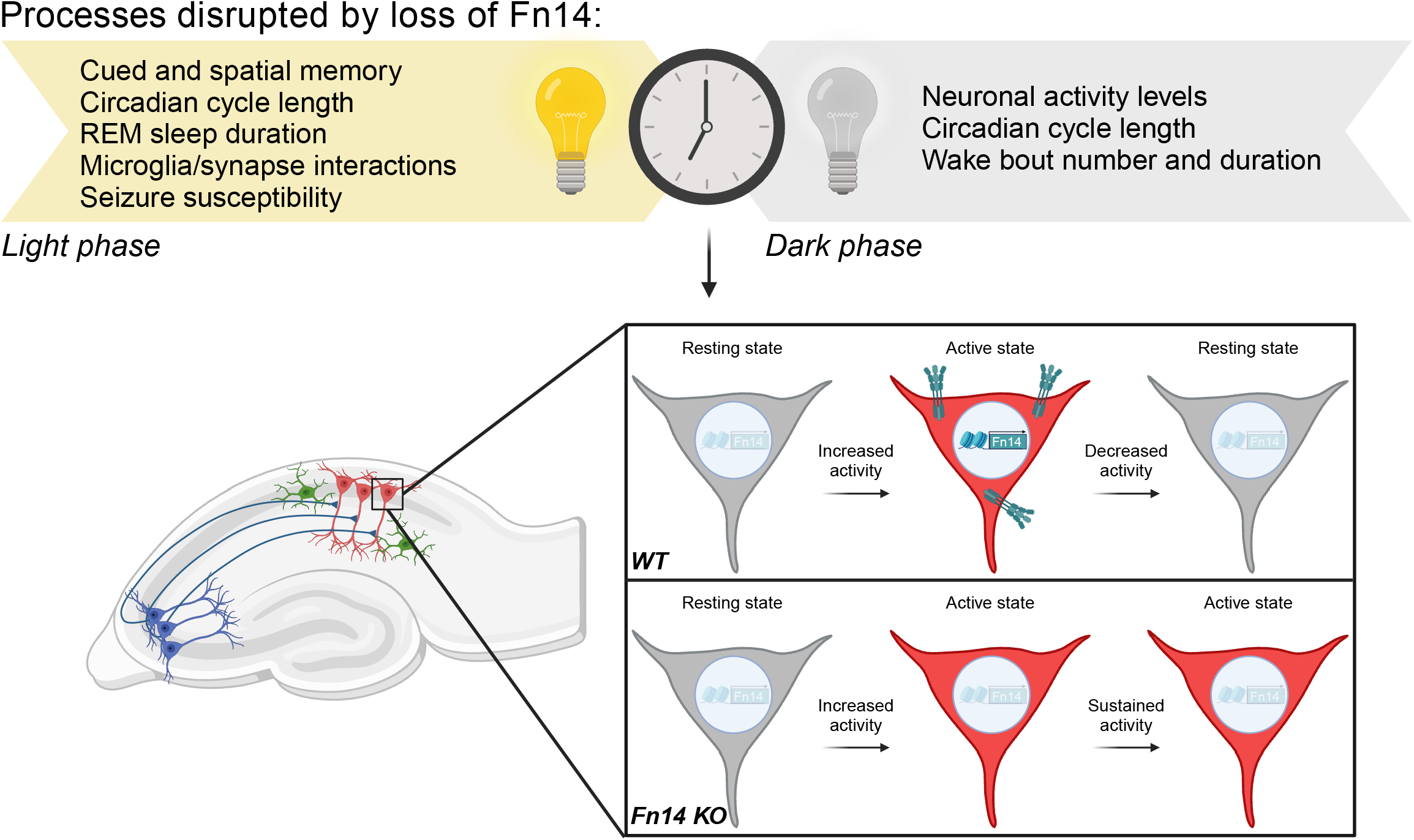
Proposed model of Fn14 function in the brain. We propose a model in which Fn14 is part of a molecular feedback loop that suppresses the activity of previously activated PYR neurons, likely to maintain circuit homeostasis. In the absence of Fn14, neurons are activated normally but remain active for a prolonged period of time, potentially contributing to the deficits in memory observed in the Fn14 KO mice. Notably, the functions of Fn14 within the hippocampus are time-of-day-dependent, consistent with the role for Fn14 in circadian rhythms and sleep-wake states identified in this study. Above, processes disrupted by Fn14 are noted along with the phase, light or dark, in which those deficits emerged.

While interactions between cytokine signaling and circadian rhythms remain incompletely understood, a growing body of evidence suggests that cytokine expression can be governed by the circadian clock, while the expression of cytokines can reciprocally contribute to clock entrainment. For instance, transcription factors that are critical for circadian period generation, such as the Cryptochrome Cry1, have been shown to be potent mediators of cytokine production and release^47^. Moreover, cytokines within the TNF family can manipulate circadian-related gene expression in both mouse and human cell lines^48^, and TNFα in particular alters the rhythmic expression of the circadian transcription factors Per1 and Per2 in cultured cells^49^. Consistent with glia (i.e. non-neuronal brain cells) being major producers of cytokines such as TNFα in the brain, the contributions of glia to circadian function are increasingly appreciated. These contributions are best understood from the perspective of astrocytes, which harbor their own molecular clock that oscillates in anti-phase with neurons of the SCN^50^. These astrocyte-specific transcriptional oscillations shape rhythmic neuronal firing and regulate the sleep-wake cycle *in vivo*^51–53^. Intriguingly, recent studies have begun to uncover roles for other populations of glia, such as oligodendroglial cells^54^ and microglia^55,56^, in mediating circadian functions in mice, although it should be noted that some features of circadian rhythms appear to remain intact in the absence of microglia^57,58^. For the most part, these data are in line with our results suggesting that Fn14 plays a role in circadian function, potentially downstream of its microglia-derived ligand TWEAK.

In combination with prior work demonstrating a role for Fn14 in sensory-dependent synapse refinement, the newly discovered role of Fn14 in circadian function suggests that this receptor may contribute to the integration of intrinsic and extrinsic influences on the brain. How might this occur? One possibility is that Fn14 is a regulator of the circadian clock within the SCN, the endogenous rhythmic pacemaker of the brain^59^. If so, the changes in circadian function observed in Fn14 KO mice could indirectly lead to impairments in neuronal activity patterns and in the functional output of neurons outside of, but connected to, the SCN, such as the hippocampus. In this scenario, changes in circadian rhythms would lie upstream of the other deficits observed in Fn14 KO mice. An alternative, but not mutually exclusive, possibility is that Fn14 mediates hippocampal function in a time-of-day-dependent manner because its expression fluctuates in PYR neurons according to the time of day. Indeed, almost all cells of the body, including hippocampal neurons^60,61^, express molecules such as CLOCK and BMAL1, which function as an intrinsic circadian clock via transcriptional/translational feedback loops with a rhythm of approximately 24 hours^59,62^. Thus, increases in CA1 neuronal activity levels in mice lacking Fn14 may fluctuate across the 24-hour cycle as a result of circadian control of Fn14 expression by clock complexes in PYR cells. Regardless of the specific cellular locus of Fn14 function, a question that remains given our use of a global Fn14 KO mouse, the results reported here support a role for Fn14 in modulating essential processes in the mature brain related to circadian biology.

While, to our knowledge, this manuscript is the first to report a role for Fn14 in modulating circadian rhythms and related behaviors, it is important to note that the TWEAK-Fn14 pathway is likely not the only TNF/TNFR family pathway to play a role in the brain. In addition to work demonstrating a role for brain-specific TNFα in regulating circadian rhythms^63^, a recent study from Pollina *et al* revealed that, in addition to Fn14, five other TNFRs were also upregulated in the hippocampus following acute kainate exposure: Tnfrsf1a, Tnfrsf1b, Ltbr, Fas, and Eda2r (Fig. S1D). Thus, the regulation of hippocampal activity and function may involve members of the TNFR family beyond Fn14. Consistent with this possibility, TNFα and its receptors TNFR1 and -2, the flagship pro-inflammatory cytokine pathways of the TNF family, have been implicated in activity-dependent synaptic scaling *in vitro* and dendritic spine remodeling in the hippocampus^64–67^. Nevertheless, whether TNF pathways other than TWEAK-Fn14 mediate core behavioral outcomes in mice, such as circadian rhythms and memory, is not yet clear.

While this study is the first to implicate Fn14 in disorders related to hyperexcitation such as epilepsy, Fn14 and its ligand TWEAK have been implicated in a diversity of other diseases associated with neuroinflammation, including neuropsychiatric lupus, multiple sclerosis, Alzheimer’s disease (AD), and stroke^20,68,69^. Perhaps most relevant to this study, Nagy *et al* found that Fn14 levels are increased in the brains of individuals with AD, and that pharmacologically dampening TWEAK in hippocampal slices from a mouse model of AD improved deficits in long-term potentiation that emerged due to Amyloid-β-mediated pathology^70^. In combination with these results, our finding that Fn14 is necessary for circadian rhythms, sleep-wake balance, and memory is in line with a possible role for TWEAK and Fn14 in AD. These findings are particularly interesting given that sleep disturbances earlier in life are a strong predictor of AD risk, but for reasons that remain unclear^71^. Thus, Fn14 could represent one of the elusive mechanistic links between circadian disruption and memory deficits in AD. Another pathological context in which Fn14 appears to be highly relevant is cancer. For example, tumor-localized TWEAK-Fn14 signaling promotes cachexia, a systemic wasting syndrome that often accompanies the terminal phase of cancer and other conditions, in mice^72^. Moreover, Fn14 has been identified as a marker and potential therapeutic target for glioma, in part because it is thought to be lowly expressed and inactive in healthy brain tissue^73,74^. However, our data indicate that, at least in mice, Fn14 is essential for mature brain function outside of a pathological context. Thus, if targeting Fn14 is to become a therapeutic strategy for treating brain cancer, the assumption that Fn14 is inactive in the healthy brain deserves revisiting.

Finally, while this study provides evidence that Fn14 coordinates hippocampal activity across multiple scales, the results should be interpreted with caveats in mind. For example, the use of a global KO mouse precludes our ability to definitively assign the functions of Fn14 that we have discovered as reflecting the roles of Fn14 expressed by neurons in particular. That said, this is the most likely explanation, especially given the dynamic upregulation of Fn14 in activated PYR neurons of CA1, the same neurons that exhibit heightened activity when Fn14 is ablated. Another point that supports Fn14 acting within the hippocampus specifically is the upregulation of AP1 transcription factor signaling in the brains of Fn14 KO compared to WT mice. Since Fn14 is known to function via the activation of transcriptional mechanisms, including through the induction of intracellular cascades that mediate the expression and activation of AP1 transcription factors^20^, seeing these changes in brain tissue supports Fn14 functioning within neurons, the brain cells that express it most highly. A second caveat is the use of kainate, which activates neurons to an extent that is largely non-physiological, as a reagent to induce neuronal activity in the hippocampus. Nevertheless, our fiber photometry data indicating that Fn14 constrains neuronal activity during normal home cage behavior validates that Fn14 functions to constrain activity even in the absence of exposure to convulsants. Despite these caveats, this study provides compelling evidence of a role for Fn14, and potentially for microglia, in a spectrum of neurological functions in healthy adult mice, ranging from the constraint of neuronal activity to the modulation of circadian rhythms.

## Acknowledgements

We thank the following individuals for critical input and feedback on the manuscript: Dr. Steve Shea (CSHL), Dr. Cliff Harpole (CSHL), Dr. Jessica Tollkuhn (CSHL), Dr. Linda Burkly (formerly Biogen), Dr. Marty Yang (UCSF), Dr. Lisa Boxer (NIH), Dr. Sameer Dhamne (Boston Children’s Hospital), Dr. Jacqueline Barker (Drexel University), Dr. Timothy Cherry (Seattle Children’s Hospital), Dr. Elizabeth Pollina (Washington University in St. Louis), Dr. Marija Cvetanovic (University of Minnesota), and Dr. Cody Walters (Nature Communications). Work in this study was aided by the Behavior and Neurophysiology Core at Boston Children’s Hospital. We also thank Biogen for providing Fn14 KO and WT mice. This work was supported by the following grants: R00MH120051, DP2MH132943, R01NS131486, Rita Allen Scholar Award, McKnight Scholar Award, Klingenstein-Simons Fellowship Award in Neuroscience, and a Brain and Behavior Foundation NARSAD grant (to L.C.); 2020 Breast Cancer Research Foundation – AACR NextGen Grant for Transformative Cancer Research (20-20-26-BORN), DoD Idea Development Grant (W81XWH2210871), and NIH Cancer Center support grant (5P30CA045508-36)(to J.C.B.).

## Author contributions

L.C. conceptualized the study. Fiber photometry and wireless telemetry experiments were designed, performed, and analyzed by A.F., L.B., A.B., A.G., and J.C.B. All other experiments were designed, performed, and analyzed by L.C., A.F., A.A., T.S., and U.V.

L.C. and A.F. wrote the first draft of the paper, which was later modified in response to input from all authors.

## STAR Methods

### Animal models

All experiments were performed in compliance with protocols approved by the Institutional Animal Care and Use Committee (IACUC) at Harvard Medical School and Cold Spring Harbor Laboratory. The following mouse lines were used in the study: C57Bl/6J (the Jackson Laboratory, JAX:000664) and B6.Tnfrsf12a^tm1(KO)Biogen^ (Fn14 KO and WT littermates^33^). Fn14 KO mice were generously provided by Dr. Linda Burkly at Biogen (Cambridge, MA) and are subject to a Material Transfer Agreement with Cold Spring Harbor Laboratory. Analyses were performed on both male and female mice between one and six months of age. No sex differences were observed in the study.

### Single-molecule fluorescence in situ hybridization (smFISH)

Sagittal or coronal sections of 20-25 μm thickness centered on the hippocampus were made using a Leica CM3050S cryostat and collected on Superfrost Plus slides then stored at -80°C until use. Multiplexed smFISH was performed using the RNAscope platform (Advanced Cell Diagnostics [ACD], Biotechne) according to the manufacturer’s instructions for fresh-frozen (multiplexed kit v1, now discontinued) or fixed-frozen (multiplexed kit v2) samples. Probes against the following transcripts were utilized: *Fn14* (*Tnfrsf12a*), *Slc17a7* (*Vglut1*), *Gad1*, *Camk2a*, *Gad2*, and *Fos*. For the quantification of *Fn14*, *Slc17a7*, and *Gad1* transcripts per cell, 60X confocal images were acquired using a LSM 710 Zeiss microscope. A total of 3 mice per condition and a minimum of two images per mouse were analyzed. *Fn14* expression was quantified using an ImageJ macro built in-house (code: www.cheadlelab.com/tools). Briefly, the DAPI channel was thresholded and binarized, and subsequently expanded using the dilate function. This expanded DAPI mask was then passed through a watershed filter to ensure that cells that were proximal to each other were separated. This DAPI mask was then used to create cell-specific ROIs, where each ROI was considered a single cell. Using these cell-masked ROIs, the number of mRNA puncta were counted using the 3D image counter function within imageJ. ROIs were classified with the following criteria: ROIs containing 3 or more *Fn14* molecules were considered positive for *Fn14*, while those containing 5 or more molecules of either *Gad1* or *Vglut1* were considered positive for each marker, respectively.

For the quantification of *Fn14*, *Camk2a*, *Gad2*, and *Fos* in mice treated with kainate or vehicle control, 40X confocal images of the hippocampus were acquired using an LSM 780 Zeiss microscope. Six images were taken per section across 4 mice per condition (kainate or vehicle [i.e. water]). Data analysis was conducted using the image processing software ImageJ (FIJI). First, a binary mask was created for the DAPI channel in each image by applying a Gaussian blur, binarizing the image, closing holes in the nuclear signal, dilating the image, and removing cells in the image that are not part of the CA1 area of the hippocampus. Using this binary mask for region of interest (ROI) analysis, the area and mean intensity were collected for each nucleus (cell) and exported into an Excel file. To calculate intensity thresholds used to determine if a cell was positive or negative for a particular marker, a supervised analysis was conducted. In this method, ten ROIs were manually selected based on if they were visually either positive or negative for a marker and their intensity values were calculated. *Gad2*, *Camk2a*, and *Fos* thresholds were extracted by collecting the mean and standard deviation of expression intensity for ROIs appearing negative for the marker and adding two standard deviations to the mean (*x̅* + 2*σ*) to calculate the threshold for each marker. *Fn14* intensity threshold was determined by selecting ROIs appearing positive for *Fn14* and calculating the threshold by subtracting two standard deviations of expression intensity from the mean (*x̅*-2*σ*).

Next, using the thresholds created for each marker, each ROI was established as either positive (represented with 1) or negative (represented with 0) for a marker. *Fn14* intensity in cells positive or negative for a given marker, or for *Fos*, was calculated. Next, average intensity values of *Fn14* were collected from cells positive for *Camk2a* and from cells positive for *Gad2* in order to compare *Fn14* intensity across cell types. Finally, the proportion of cells co-expressing *Fn14* and each cell type marker was calculated.

### Behavior

#### Cued Fear Conditioning

On training day, subjects were placed into a square fear-conditioning arena of 24(w)x20(d)x30(h) cm equipped with a shock grid floor and acrylic walls patterned with horizontal black and white bars 2 cm in width. Subjects were allowed to acclimate to the arena for 4 minutes before data acquisition. During training, mice were presented with three 20 second tones (75 dB; 2000 Hz) followed by a 2 second foot shock (0.5 mA) with variable inter-trial intervals totaling 5 minutes. After training, subjects were returned to their home cages for 24 hours before being tested in familiar and novel contexts. For familiar context (the paired context without the cued tone; Context (-) tone) subjects were re-acclimated to the test arena for 5 minutes without receiving tone cues or shocks to reduce freezing to non-tone cues. After testing freezing in the Context - tone condition and on the same day, subjects were exposed to a novel context (circular arena 30(w) x 30(h) cm, with clear acrylic floor and polka-dot walls) for 3 minutes to habituate the mice to the novel context before freezing was measured. Mice were then returned to their home cages for 24 hours before being re-exposed to the novel context, then they were re-exposed to the cued tone (75 dB; 2000 Hz) for three minutes during acquisition. Freezing was calculated using Ethovision XT v. 15 (Nodulus, Netherlands) with activity detection set to 300 ms, and data were presented as freezing over the trial time.

#### Morris Water Maze

Each training trial consisted of four 90 s sub-trials in which each subject’s starting position was pseudo-randomized to each of the four cardinal directions in a 137 cm wide water bath containing 24°C clear water filled up to 25 cm from the rim of the tub. The cardinal directions were marked on the wall of the tub with 20 cm diameter symbols. Subjects were initially trained over two trials where the goal zone was visible (visible trials), where the goal platform was raised 0.5 cm above the water line and was marked with a bright flag for increased visibility. Each trial ended either after the trial time expired, or after the subject correctly found and stayed on the goal platform for more than 5 seconds. If a mouse did not find the platform within 90 seconds, it was gently moved to the platform and left there for 5 seconds. The day following visible platform training, the goal platform was submerged (0.5 cm below the water line) and moved to a different quadrant. Subjects were tested on the hidden platform over 5 consecutive trials spanning 48 hours. On the fourth day (probe trial) the goal platform was removed from the testing arena and subjects were placed facing the wall opposite of the previous goal platform’s position. Subjects were allowed to swim for a total of 60 s before being removed from the arena. On reversal trials (4 trials), the goal platform remained submerged, but was moved to the opposite end of the arena. Subjects started the reverse trials facing the furthest wall and were allowed to search for the goal platform for 90 s. If the subject failed to find the goal platform, the subject was oriented in the correct direction and guided to the goal platform before being removed from the arena. Latency to goal platform, distance swam, and subject position were collected using Ethovision XT v. 15 (Nodulus, Netherlands).

#### Optomotor testing of visual acuity

An optomotor device (CerebralMechanics, Canada) was used to measure visual acuity. The apparatus consists of 4 computer monitors arranged in a square, in order to produce a virtual 3-D environment, with a lid to enclose subjects. Using the Optomotor 1.7.7 program, a virtual cylindrical space with vertical sinusoidal gratings was drawn on the monitors such that each monitor acted as a virtual window into the surrounding cylindrical space. Mice were placed on a lifted platform in the optomotor device and allowed to move freely, and tracking software was used to position the center of the virtual cylinder at the mouse’s head. Typically, when the cylinder with the grating stimuli is rotated (12 deg/sec), mice will begin to track the grating stimuli across the virtual space with reflexive head movements in concert with the stimulus motion. If the mouse’s head tracked the cylinder rotation, it was judged that the animal could see the grating. Using a staircase procedure, the mouse was tested systematically against increasing spatial frequencies of the grating until the animal no longer responded, with the mouse’s acuity being assigned as the highest spatial frequency that the mouse responded to by tracking.

### Fiber photometry

#### Stereotaxic Surgery (Viral Injections and Optic Fiber implants)

All surgical procedures were performed in line with CSHL guidelines for aseptic technique and in accordance with the humane treatment of animals as specified by the IACUC. At the start of surgical procedures, mice were anesthetized with isoflurane (3% induction; Somnosuite, Kent Scientific), and then injected with buprenorphine SR (Zoopharm, 0.5 mg/kg, s.c.). Upon confirmation of deep anesthesia mice were placed into a stereotaxic frame (David Kopf Instruments) where they were maintained at 1-1.5% isoflurane. A midline incision was then made from the posterior margin of the eyes to the scapulae to expose the braincase. The skull was cleaned and then a drill was positioned over the skull to drill a hole for the viral injection. Mice were then injected unilaterally within the dorsal hippocampus (-2.06 mm AP, 1.3 mm ML, 1.25 mm DV) using a 30-gauge blunt Neuros syringe (Hamilton) at a rate of 20 nl/min for a total volume of 200 nL. AAV9-CamkII-GCaMP6f (viral titer 1x10^13^ gc/mL), obtained from Addgene, was injected. After the infusion, the needle was left in place for at least 10 minutes before the microinjector (World Precision Instruments) was withdrawn slowly. Directly following virus injection, a fiber optic (400 um in diameter; 0.48 NA, Doric Lenses) was lowered just dorsally to the injection site (-2.06 mm AP, 1.3 mm ML, 1.20 mm DV). The optic implant was then fixed in place with Metabond (Parkell) and dental cement. After surgery, mice were then allowed to recover until ambulatory on a heated pad, then returned to their home cage with Hydrogel and DietGel available. Mice were then allowed to recover for approximately 4 weeks to allow for viral expression before behavioral experiments and fiber photometry recordings began.

#### In vivo optical recording

Approximately 4 weeks after viral transduction and fiber optic implantation, baseline recording sessions began. In brief, mice were tethered to a fiber optic patch cord (400 uM, Doric Lenses) via a ceramic mating sleeve connected to the implanted optic fiber (400 uM, Doric Lenses), and fiber photometry data was collected using a fiber photometry setup with optical components from Doric Lenses and controlled by a real-time processor from Tucker Davis Technologies (TDT; RZ5P). TDT Synapse software was used for data acquisition, where LED sources of 465 nm (Signal / GCaMP) and 405 nm (Control / Isosbestic) were modulated at 211 or 230 Hz and 330 Hz, respectively. LED currents were adjusted in order to return a voltage between 100 and 150 mV for each signal, and were offset by 5 mA. The signals were then demodulated using a 6 Hz low-pass frequency filter, where subsequent analysis occurred. In brief, GuPPy, an open-source Python toolbox for fiber photometry analysis^75^, was used to compute ΔF/F and z-score values, as well as Ca^2+^ event amplitude and frequency, for all recordings. We did not analyze the first minute of each 11-minute epoch in order to remove any artifacts that may occur as the recording begins (i.e. 600 seconds was analyzed for each epoch). To calculate the change in fluorescence ΔF/F from the photometry signal F, GuPPy normalized the data by fitting the GCaMP6f signal with the isosbestic control wavelength and computing ΔF/F = Signal - Fitted Control. It then computed a standard z-score signal for the ΔF/F data using z score=F/F-(mean of F/F) standard deviation of F/F to evaluate the deviation of the ΔF/F signal from its mean. We incorporated a 600-second user-defined window for thresholding calcium transients in the ΔF/F and z-score traces; GuPPy identifies the average amplitude and frequency (defined as events per minute) of the transients in each trace, as well as the amplitude and timing of each transient. Transients were identified by filtering out events with amplitudes greater than two times the median absolute deviation (MAD) above the median of the user-defined window and finding peaks greater than three MADs above the resulting trace. We identified the maximum z-score amplitude for each epoch by finding the largest amplitude in the table of transient timestamps and amplitudes outputted by GuPPy. We used a custom Jupyter Notebook script to calculate area-under-the-curve (AUC) for ΔF/F and z-score traces in 10-minute time bins. A MATLAB script was additionally used to determine the average amplitudes of all values, all positive values, and all absolute values for the ΔF/F and z-score traces.

### Quantification of AP-1 activation

Whole brain tissue was collected from Fn14 KO and WT mice at P27 and flash frozen in liquid nitrogen. Tissue was later thawed and homogenized in RIPA buffer (VWR) via agitation on ice for 30 minutes before centrifugation at 23,000 x *g* for 10 minutes. 5 microliters of the insoluble fraction were then diluted in Complete Lysis Buffer (Active Motif) and nuclear protein concentration was determined using a Bradford assay (Bio-rad). Once nuclear proteins were diluted to equal concentrations in Complete Lysis Buffer, 20 μg of sample was then used to quantify binding of Fos and phosphorylated Jun (P-Jun) to oligonucleotide consensus binding sites for AP-1 family members according to the manufacturer’s instructions. Briefly, nuclear extracts were added to a pre-coated 96-well plate, and antibodies against P-Jun and Fos were added and the plate was incubated for 1 hour at room temperature. After washing each well, an HRP-conjugated secondary antibody against P-Jun or Fos was added and the plate was incubated at room temperature for another hour. After washing off the unbound secondary antibody, each colorimetric reaction was developed and subsequently stopped using Stop solution. Absorbance at 450 nm was measured for protein binding within 5 minutes of addition of Stop solution with 650 nm as a reference. Technical replicates (n = 2/sample) were averaged and data was normalized to WT samples.

### RNA isolation and rt-qPCR

Fn14 KO and WT littermate mice at P27 were euthanized and their brains were bisected and flash frozen using liquid nitrogen in 1 mL of Trizol (Ambion) and kept at - 80°C until processing. Tissue was then homogenized using a motorized tissue homogenizer (Fisher Scientific) in a clean, RNAase-free environment. Once homogenized, 200 μL of chloroform was added to each sample and, after thorough mixing, samples were centrifuged at 21,000x*g* for 15 minutes for phase separation. The colorless phase was then collected and combined with equal volume of 70% ethanol and used as input in the RNeasy Micro kit (Qiagen), after which the manufacturer’s instructions were followed to further purify the RNA. RNA concentration was then determined using a nanodrop (ND 1000; NanoDrop Technologies Inc), and once RNA samples were diluted to equal concentrations, samples were converted into cDNA using SuperScript™ III First-Strand Synthesis System (Thermo Fisher) following the manufacturer’s instructions. The transcript encoding Scn1a was then amplified and detected using Power Up Sybr Green (Thermo Fisher) in a Quant Studio 3 Real-Time PCR system (Thermo Fisher). Crossing threshold (Ct) values were calculated using the QuantStudio program and relative expression, 2^-ΔΔCt^, was calculated using *GAPDH* as a reference control.

### Circadian rhythms

Mice 2-3 months of age were separated and singly housed in conventional cages with the addition of wireless running wheels (Med Associates Inc: ENV-047). Mice were allowed to acclimate to their respective running wheels for 3-5 days before data acquisition. After acclimation, activity was recorded by measuring the number of running wheel rotations every minute using a wireless recording hub and associated software (Med Associates Inc: DIG-807, SOF-860). Mice were kept in normal environmental conditions within the vivarium, which is kept on a 12-hour:12-hour light/dark cycle, for 10-14 days before being placed into constant darkness for an additional 10-14 days of acquisition. After acquisition of their running wheel activity in both the 12:12 light/dark cycle and constant darkness (to record free running activity), running wheel data was parsed into these environmental conditions: 12:12 LD and constant darkness. Both datasets were then analyzed using a custom MatLab script which, in short, normalized the running wheel activity within a given mouse to the mouse’s mean running activity, and then iteratively fit sinusoidal waves to the data to find the wave with the best fit to the activity data. The period of this resultant sinusoid function was then reported as the running wheel activity period of a given mouse.

### Wireless telemetry (sleep-wake dynamics)

Mice were deeply anesthetized under isoflurane vapors (3% induction, 1.5% maintenance) and implanted with HD-X02 biotelemetry transmitters (Data Sciences International, DSI, St. Paul, MN, USA) to allow acquisition of electroencephalogram (EEG) and electromyogram (EMG) potentials. Following immobilization in a stereotaxic apparatus, a midline incision was made extending between the caudal margin of the eyes and the midpoint of the scapulae. The skull was exposed and cleaned, and two stainless steel screws (00-96 x 1/16; Plastics One, Roanoke VA, USA) were inserted through the skull to make contact with the underlying dura mater. These screws served as cortical electrodes. One screw was placed 1 mm lateral to the sagittal suture and 1 mm rostral to Bregma. The other screw was placed contralaterally 2 mm from the sagittal suture and 2 mm caudal to Bregma. The transmitter was inserted into a subcutaneous pocket along the back of the animal. A set of leads was attached to the cortical electrodes and secured with dental cement. Another set of leads was inserted and sutured into the trapezius muscles for EMG measurement. The surgical procedures were performed using aseptic technique, and buprenorphine SR (0.05 mg/kg, SC) was administered to provide post-operative analgesia along with supplemental warmth (heating pad) until the animals were mobile. Following surgery, mice were singly housed and their cages were placed on top of receiver boards (RPC-1; DSI). These boards relay telemetry data to a data exchange matrix (DSI) and a computer running Ponemah software (version 6.1; DSI, St. Paul, MN, USA). Mice were allowed to recover from the surgery for 2 weeks prior to beginning sleep recordings.

For analysis, raw biopotentials were band pass filtered (0.3-50 Hz for EEG, and 10-100 Hz for EMG) and analyzed in 5 second epochs as previously described^40^. The delta band was set at 0.5–4.0 Hz, and the theta band was set at 6-9 Hz. Artifact detection thresholds were set at 0.4 mV for both EMG and EEG, and if >10% of an epoch fell outside this threshold, the entire epoch was scored as artifact. Wake was characterized by high frequency and low voltage EEG accompanied by high voltage EMG. NREM (i.e., slow wave sleep) sleep was characterized by low frequency and high voltage EEG (predominant delta), accompanied by low voltage EMG. REM (i.e., paradoxical) sleep was characterized by high frequency, low voltage EEG (predominantly theta) and EMG values. Five second epochs were collapsed into 1-hour bins for subsequent graphing and statistical analyses. For spectral analyses, biopotentials were visually inspected, cleaned of artifacts, and subjected to Fast-Fourier transforms. Periodogram data were collected in 5-second epochs of scored data and then the EEG power spectra for each vigilance state was compared between genotypes and at different times of day.

### Wireless Telemetry (baseline and sleep rebound recordings)

Mice were given a 24-hour acclimation period before telemetry was used to obtain EEG, EMG, body temperature, and locomotor activity continuously for 48 hours. During the first 24 hours, baseline sleep and wake data were collected and the mice were undisturbed. At the start of the next light cycle (ZT0-ZT6), mice were sleep-deprived by gentle handling for six hours^40^. Recovery sleep and wake data were then recorded over the subsequent 18 hours. All data were processed and analyzed using DSI Neuroscore software. Baseline and recovery recordings were scored as either wake, non-rapid eye movement (NREM) or rapid eye movement (REM) sleep in 5-second bins. Scorings were then analyzed in 1-hour bins for number of bouts, average bout length, and percent coverage of each sleep stage. Baseline and recovery EEG recordings were also automatically analyzed using Neuroscore for delta, theta, gamma and alpha spectral power; power density (amplitude); transitions between sleep stages; and number of microwakes (wake bouts of less than 5 seconds in duration).

### Immunofluorescence

WT and Fn14 KO mice were perfused with ice cold PBS (Gibco) and 4% for paraformaldehyde (PFA), then the whole brains were harvested and post-fixed for 12 hours. After fixation, tissue was incubated in 15% and then 30% sucrose solution before being embedded in OCT (-80°C). Embedded tissue was sectioned coronally at 25 μm thickness onto Superfrost Plus slides using a Leica CM3050 S cryostat. Sections were then washed in PBS and blocked in blocking solution (PBS adjusted to 5% normal goat serum [NGS] and 0.3% Triton X-100 [TX-100]) for 1 hour at room temperature before being incubated in primary antibody solution containing Chicken anti-Iba1 (Synaptic Systems, 234 009; [1:1000]), Rabbit-anti-Vglut1 (Invitrogen YA364832 [1:1000]), and Mouse-anti-Vgat (Synaptic Systems, 131 001; [1:1000]) antibody diluted in PBS with 5% NGS and 0.1% TX-100 (probing solution), overnight at 4°C. The next day, sections were washed 3 times for 10 minutes per wash in PBS before incubation in secondary antibodies Alexafluor 488 goat anti-rabbit (Abcam 150077; [1:500]) Alexafluor 555 rabbit anti-goat (Thermofisher A21428; [1:1000]) and Alexafluor 488 chicken anti-rabbit (synaptic systems160 026; [1:1000]) diluted in probing solution for 2 hours at room temperature. Sections were then washed in PBS, covered with DAPI fluoromount-G (SouthernBiotech), and cover-slipped.

#### Microglia-Synapse Interactions

Z-stack images (40X, numerical aperture 1.4) were obtained on a confocal (LSM 780 Zeiss) Microscope. Two sections per mouse (n = 3-4 mice/genotype) containing CA1 were imaged as a Z-stack (3008 x 3008 pixels, voxel = 70.7 x 70.7 x 311 nm [x,y,z], 16-bit). Images were then converted from .CZI to .IMS files to quantify in Imaris 10.0.0, using the Imaris File Converter. A background subtraction (53.1 µm) and gaussian filter (0.0707 µm) were applied to all images under image processing in this program. Representative 3-dimensional surfaces of microglia (Iba1), Vglut1, and Vgat signals were then reconstructed in Imaris. In brief, surfaces were created using a signal intensity threshold based on the average signal intensity of a given object within the imaging field. After surfaces were created, relative distances between objects were determined and Vglut1 and Vgat puncta were then filtered and classified as being within -0.07 and 0.07 µm from a microglial surface. The stringent distance-based filter allowed us to filter out synaptic puncta that are more likely to reside within the glial cell (i.e. to have been engulfed by the cell) rather than in contact with the surface of the cell. Average values of volume and number of surface objects, denoted under “sum”, “mean”, and “count,” were exported for statistical analysis.

### EEG recordings and PTZ seizure induction

#### EEG telemetry unit implantation

Mice were implanted with wireless telemetry units (PhysioTel ETA-F10; DSI, Data Sciences International) under sterile techniques per laboratory protocol as described above. Under anesthesia, a transmitter was placed intraperitoneally, and electrodes were threaded subcutaneously to the cranium. After skull exposure, haemostasis, and identification of the cranial sutures bregma and lambda, two 1-mm diameter burr holes were drilled over the right olfactory bulb (reference) and left occipital cortex (active). The epidural electrodes of the telemetry units, connected to the leads of the transmitter, were placed into the burr holes, and secured using stainless steel skull screws. Once in place, the skull screws were covered with dental cement. Mice were subcutaneously injected 0 and 24 hours post-operatively with 5 mg/kg meloxicam for analgesia. After 1 week of recovery, mice were individually housed in their home cages in a 12/12 light/dark cycle, within a temperature- and humidity-controlled chamber with *ad libitum* access to food and water.

#### Baseline and PTZ seizure induction

After a 24-hour acclimation period, one-channel EEG was recorded differentially between the reference (right olfactory bulb) and active (left occipital lobe) electrodes using the Ponemah acquisition platform (DSI). EEG, core-body temperature, and locomotor activity signals were continuously sampled from all mice for 48 hours along with time-registered videos. At the end of baseline acquisition, all mice were provoked with a convulsive dose (60 mg/kg; i.p.) of the GABA_a_ receptor antagonist pentylenetetrazole (PTZ; Sigma-Aldrich, Co.) to measure seizure susceptibility and evaluate seizure thresholds^46,76–78^. Mice were continuously monitored for clinical and electrographic seizure activity for 20 minutes.

#### Data analysis

All data were processed and analyzed using Neuroscore software (DSI). Baseline EEG was analyzed for spontaneous seizure activity, circadian biometrics, and spectral power band analysis^76,77^. Relative spectral power in delta (1-4 Hz), theta (4-8 Hz), alpha (8-12 Hz), beta (12-30 Hz), low gamma (30-60 Hz) and high gamma (60-90 Hz) frequency bands of the baseline EEG were calculated using the fast Fourier transform (FFT) technique.

PTZ-induced seizure activity was broadly scored on a modified Raccine’s scale as electrographic spikes (score: 1), myoclonic seizures (score: 3), generalized tonic-clonic seizures (GTC; score: 5) and death (score: 6). Per mouse, number of myoclonic seizures, latency and incidence of GTC seizures, number of GTCs, and total duration of GTC were recorded. Mice without seizures were assigned a time of 20 min at the end of the PTZ challenge observation period.

### Statistical analyses

For all analyses, sample sizes were chosen based on previously generated data. Acquired data was first tested for normality and log-normality before choosing a parametric or non-parametric statistical test. When the data were found to be normal, parametric t-tests, one-way ANOVAs, or repeated measures two-way ANOVAs were used. If data was found to be non-gaussian and non-logarithmic, a Mann-Whitney test was performed.

Statistical analyses were performed in Excel and Prism 9.0 (GraphPad Software). Figures were created using MATLAB R2019b and Graphpad Prism and formatted using Adobe Illustrator (2024). The model in Figure 7 was generated in biorender.com. Data are presented as mean ± SEM unless otherwise indicated.

