## Supplemental figures and legends for "The cytokine receptor Fn14 is a molecular brake on neuronal activity that mediates circadian function *in vivo*"

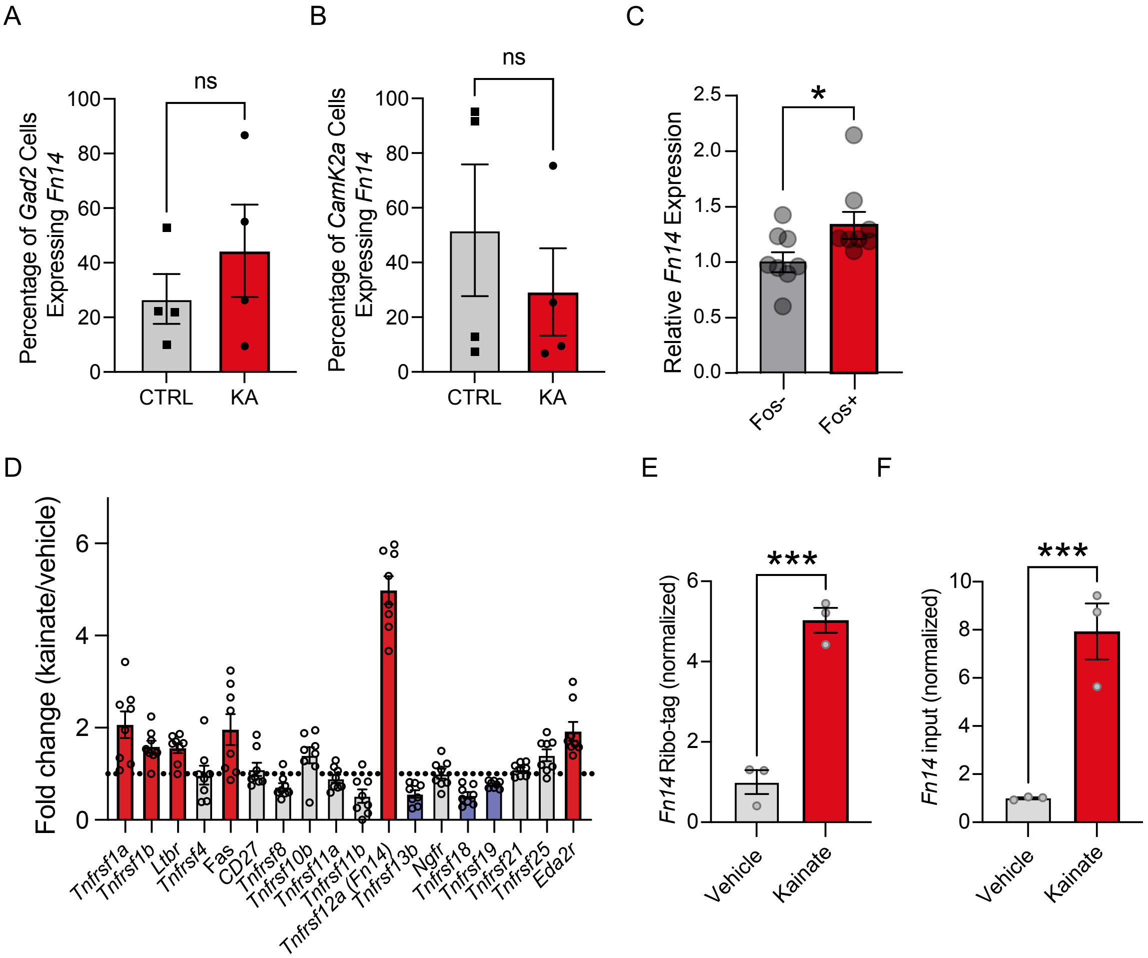


Figure S1 (Related to Figure 1). Activity-dependent expression of Fn14 in the hippocampus.

**Figure S1 (Related to Figure 1). Activity-dependent expression of Fn14 in the hippocampus.** (A),(B) Quantification of the percentage of *Gad2*+ inhibitory neurons (A) and *Camk2a*+ PYR neurons (B) that express *Fn14* in response to kainate. Unpaired Student's T-tests, $p > 0.05$. (C) Relative Fn14 expression in Fos- and Fos+ cells aggregated across control and kainate- treated mice (unpaired Student's T-Test, *$p < 0.05$). (D) Fold change in expression of genes encoding Tumor Necrosis Factor Receptor Superfamily members in the hippocampus after exposing mice to kainate for two hours, data re-analyzed from Pollina et al, 2023. Dashed line = 1 (no change). Red bars, genes that were significantly upregulated by kainate exposure; blue bars, genes that were downregulated by kainate exposure; gray, genes that were unchanged by kainate exposure. (E) Normalized quantification of Fn14 expression in ribosomal RNA extracted from PYR neurons, re-analyzed from data in Yap et al, 2023. Unpaired Student's t test, ***$p < 0.001$. (F) Normalized quantification of Fn14 expression in the input fraction from the same experiment. Unpaired Student's T-Test, ***$p<0.001$.

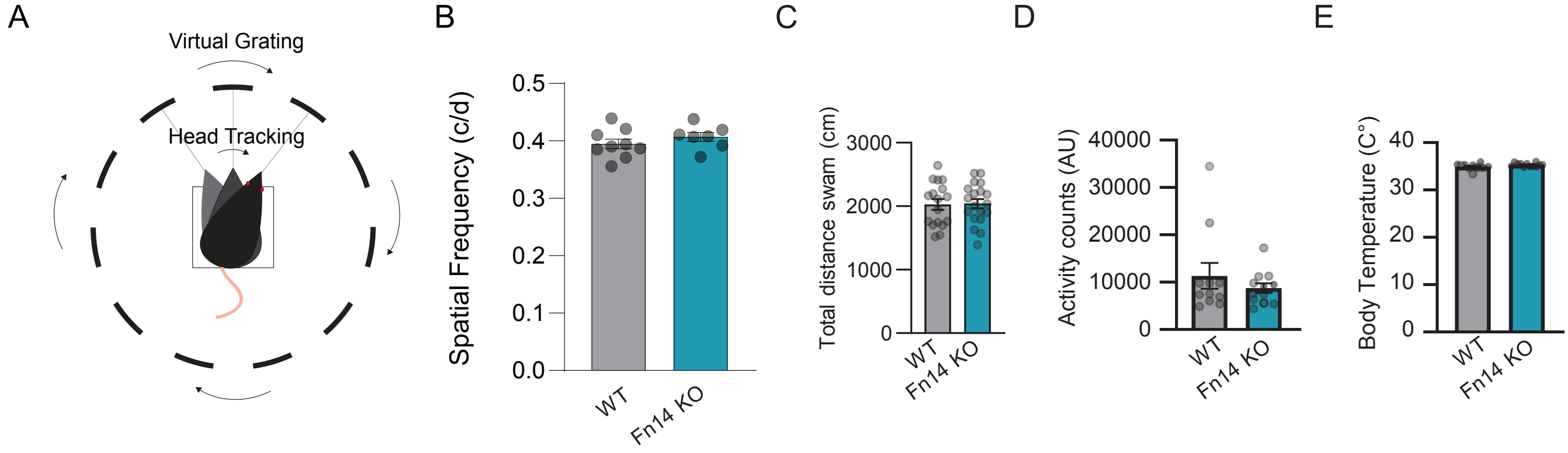


Figure S2 (Related to Figure 2). Visual acuity and locomotor activity are normal in the absence of Fn14.

**Figure S2 (Related to Figure 2). Visual acuity and locomotor activity are normal in the absence of Fn14**. (A) Schematic of the optomotor task in which four monitors surrounding the mouse create a virtual visual grating with varying spatial frequency. When the grating is perceptible, the mouse will track the grating with a stereotyped head movement. (B) Quantification of visual acuity using unpaired Student's T-test between WT ($n = 9$) and Fn14 KO mice ($n = 7$), $p > 0.05$. (C) Total distance swam by mice during the Morris Water Maze probe trial. (D) Overall activity levels in Fn14 KO and WT mice measured by automated detection in video recordings during the EEG experiments. (E) Body temperatures were largely equivalent in Fn14 KO and WT mice. For (C) – (E) Unpaired Student's T tests, $p > 0.05$.

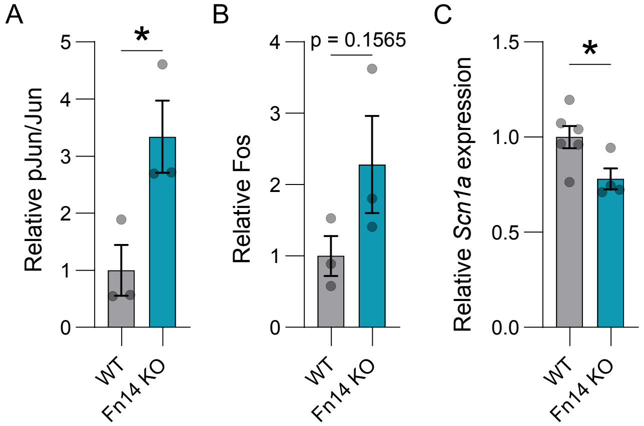


Figure S3 (Related to Figure 3). AP1 transcription factor activity is increased in the brains of Fn14 KO mice.

**Figure S3 (Related to Figure 3). AP1 transcription factor activity is increased in the brains of Fn14 KO mice**. (A) ELISA quantification of the relative amount of phosphorylated Jun versus un-phosphorylated Jun in brain homogenates normalized to WT (n = 3 mice/genotype; unpaired Student's T-test: *$p<0.05$). (B) ELISA quantification of relative Fos protein concentration in brain homogenates normalized to WT (n = 3 mice/genotype; unpaired Student's T-test: $p = 0.1565$). (C) RT-qPCR quantification of relative *Scn1a* expression in brain homogenates normalized to GAPDH and plotted as normalized to WT; n = 4 mice/genotype; unpaired Student's T-Test: *$p < 0.05$.

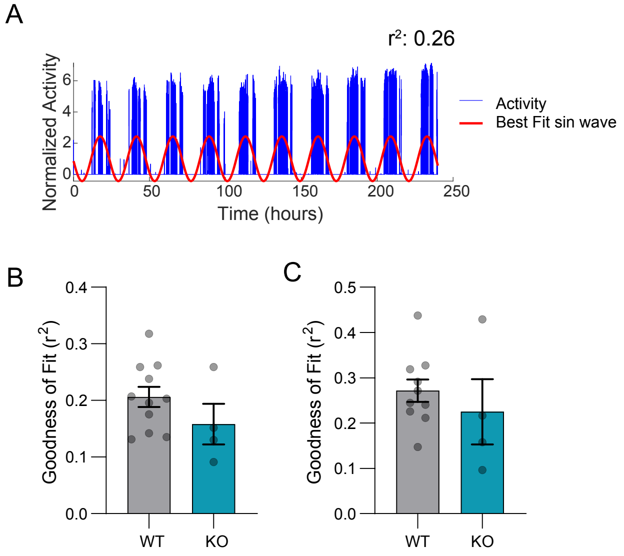


Figure S4 (Related to Figure 4). Goodness of fit of the sin wave in the analysis of running wheel data.

**Figure S4 (Related to Figure 4). Goodness of fit of the sin wave in the analysis of running wheel data**. (A) Example normalized activity data (blue) from an individual mouse over a 10-day period of time with the best fit sin wave shown in red. (B) Non-linear regression goodness of fit of WT and Fn14 KO mice in 12:12 light/dark conditions (Welch's t-test, p = 0.2883). (C) Non-linear regression goodness of fit of WT and Fn14 KO mice in constant dark (Welch's t-test, p = Welch's t-test, p = 0.5775).

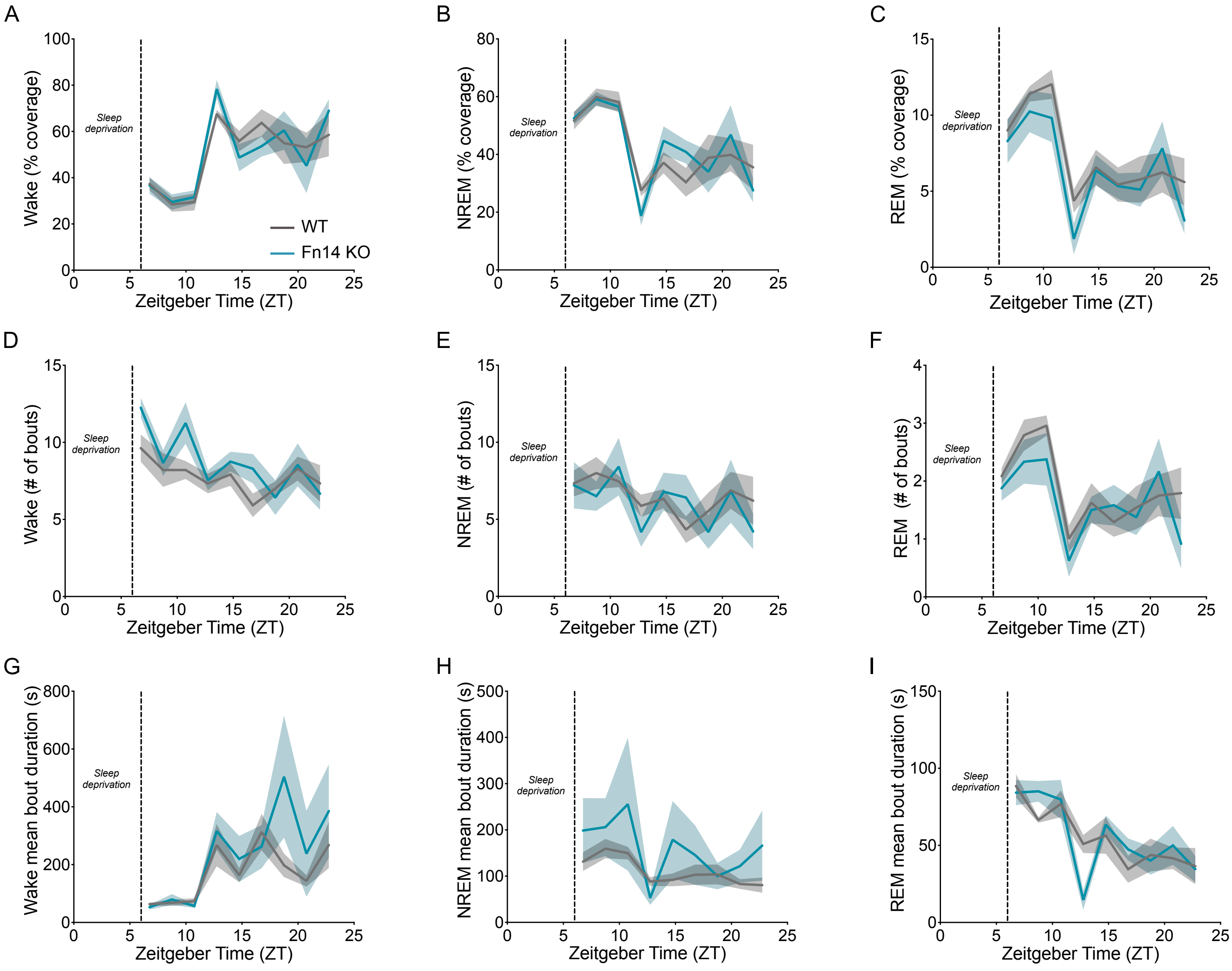


Figure S5 (Related to Figure 4). Patterns of sleep recovery following sleep deprivation in Fn14 KO and WT mice.

**Figure S5 (Related to Figure 4). Patterns of sleep recovery following sleep deprivation in Fn14 KO and WT mice**. (A) – (C) Percent coverage of wake (A), NREM sleep (B), and REM sleep (C) in Fn14 KO and WT mice. (D) – (F) number of wake (D), NREM sleep (E), and REM sleep (F) bouts in Fn14 KO and WT mice. (G) – (I) bout duration for wake (G), NREM sleep (H), and REM sleep (I) in Fn14 KO and WT mice.

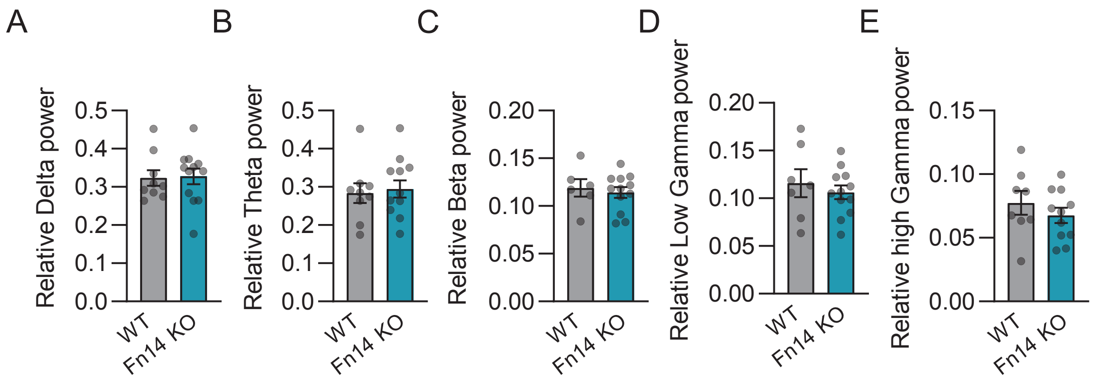


Figure S6 (Related to Figure 6). Averaged macroscopic brain activity is largely normal in the Fn14 KO.

**Figure S6 (Related to Figure 6). Averaged macroscopic brain activity is largely normal in the Fn14 KO.** (A) – (E) Averaged EEG spectral power across frequency bands were equal between Fn14 KO and WT mice. Frequency shown on y axis. ($n = 11$ WT and 12 KO mice, unpaired Student's t-tests, $p > 0.05$).
